# Inferring genome-wide correlations of mutation fitness effects between populations

**DOI:** 10.1101/703918

**Authors:** Xin Huang, Alyssa Lyn Fortier, Alec J. Coffman, Travis J. Struck, Megan N. Irby, Jennifer E. James, José E. Léon-Burguete, Aaron P. Ragsdale, Ryan N. Gutenkunst

## Abstract

The effect of a mutation on fitness may differ between populations depending on environmental and genetic context, but little is known about the factors that underlie such differences. To quantify genome-wide correlations in mutation fitness effects, we developed a novel concept called a joint distribution of fitness effects (DFE) between populations. We then proposed a new statistic *w* to measure the DFE correlation between populations. Using simulation, we showed that inferring the DFE correlation from the joint allele frequency spectrum is statistically precise and robust. Using population genomic data, we inferred DFE correlations of populations in humans, *Drosophila melanogaster*, and wild tomatoes. In these specices, we found that the overall correlation of the joint DFE was inversely related to genetic differentiation. In humans and *D. melanogaster*, deleterious mutations had a lower DFE correlation than tolerated mutations, indicating a complex joint DFE. Altogether, the DFE correlation can be reliably inferred, and it offers extensive insight into the genetics of population divergence.

## Introduction

New mutations that alter fitness are the key input into the evolutionary process. Typically, the majority of new mutations are deleterious or nearly neutral, and only a small minority are adaptive. These three categories constitute a continuum of fitness effects—the distribution of fitness effects (DFE) of new mutations (Eyre-Walker and Keightley 2007). The DFE is central to many theoretical evolutionary topics, such as the maintenance of genetic variation (Charlesworth 1994) and the evolution of recombination (Barton 1995), in addition to being key to applied evolutionary topics, such as the emergence of pathogens (Gandon et al. 2013) and the genetic architecture of complex disease (Durvasula and Lohmueller 2019).

The DFE can be quantified by either experimental approaches or statistical inference. Experimental approaches measure the DFE using random mutagenesis (Elena et al. 1998) or mutation accumulation (Fry et al. 2002); however, these approaches are limited to studying a small number of mutations. Most of our knowledge regarding the DFE has come from statistical inferences based on contemporary patterns of natural genetic variation. In these inferences, genetic data are typically summarized by the allele frequency spectrum (AFS). In some methods, a demographic model is inferred from the AFS of putatively neutral variants, and the DFE is estimated from the AFS of variants under selection, conditional on the best fit demographic model (Eyre-Walker et al. 2006; Keightley and Eyre-Walker 2007; Boyko et al. 2008; Kim et al. 2017). In other methods, the background pattern of variation is accounted for by the inclusion of nuisance parameters when fitting a DFE model to the AFS of variants under selection (Eyre-Walker et al. 2006; Tataru et al. 2017; Barton and Zeng 2018). In an alternative approach, a recent study applied approximate Bayesian computation to simultaneously infer the DFE and a demographic model (Johri et al. 2020). Moreover, a linear regression method can be used to infer the DFE from nucleotide diversity (James et al. 2017). These approaches has been applied to numerous organisms, including plants (Chen et al. 2017; Huber et al. 2018; Chen et al. 2020), *Drosophila melanogaster* (Keightley and Eyre-Walker 2007; Huber et al. 2017; Castellano et al. 2017; Barton and Zeng 2018; Johri et al. 2020), and primates (Boyko et al. 2008; Huber et al. 2017;Kim et al. 2017; Ma et al. 2013; Castellano et al. 2019).

Using these inference methods, several studies have found evidence for differences in DFEs among different populations (Boyko et al. 2008; Ma et al. 2013; Kim et al. 2017; Castellano et al. 2019; Tataru and Bataillon 2019). These studies, however, have been limited by the implicit assumption that the fitness effects of a given mutation in different populations are independent draws from distinct DFEs. Biologically, the fitness effects of a given mutation in different contexts are correlated, but it is unclear how large this correlation is and what factors affect it. Moreover, these studies only compared DFEs from the AFS of single populations and therefore cannot investigate differences in fitness effects in new environments after population divergence.

Here, we developed a novel concept called the joint DFE of new mutations, which can be inferred from the joint AFS of pairs of populations. We then defined the correlation of mutation fitness effects between populations using the joint DFE. With simulation, we showed that inferring the joint DFE and correlation requires only modest sample sizes and is robust to many forms of model misspecification. We then applied our approach to data from humans, D. melanogaster, and wild tomatoes. We found that the correlation of mutation fitness effects between populations is lowest in wild tomatoes and highest in humans. In *D. melanogaster* and wild tomatoes, we found differences in the correlation among genes with different functions. We also found that mutations with more deleterious effects exhibit lower correlations. Together, our results show that the joint DFE and correlation of mutation fitness effects offer new insight into the population genetics of these species.

## Results

### Definition

To define the joint DFE, we considered two populations that have recently diverged, one of which may have entered a new environment (Fig. 1A). We also considered that a mutation has selection coefficient *s*_1_ in the ancestral population and *s*_2_ in the recently diverged population. For two populations, the joint AFS is a matrix in which each entry *i, j* corresponds to the number of variants observed at frequency *i* in population 1 and *j* in population 2 in a sequenced sample of individuals from the two populations. Different combinations of *s*_1_ and *s*_2_ lead to distinct patterns in the joint AFS (Fig. 1B). We refer to the joint probability distribution for (*s*_1_, *s*_2_) as the joint DFE (Fig. 1C), and we refer to the marginal probability distributions for *s*_1_ or *s*_2_ as the marginal DFEs for population 1 or population 2, respectively. The observed AFS from a pair of populations results from integrating spectra for different values of *s*_1_ and *s*_2_ over the joint DFE.

**Figure 1:**
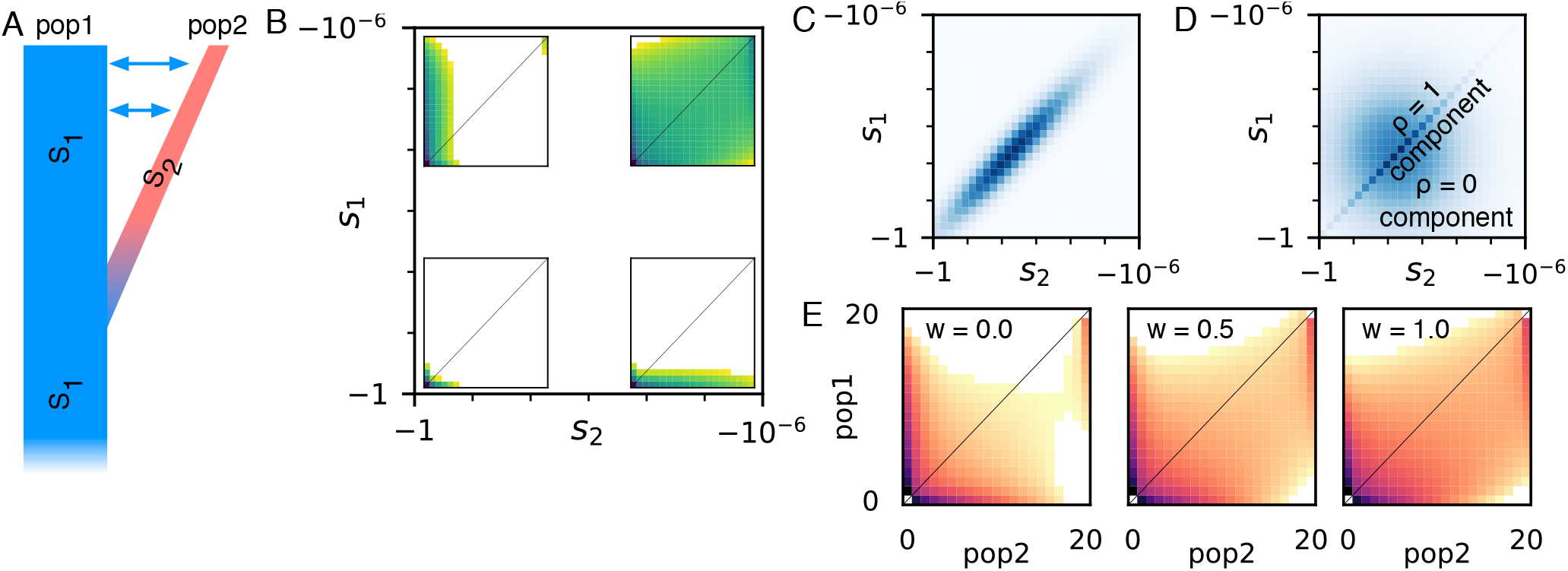
The joint allele frequency spectrum (AFS) and joint distribution of fitness effects (DFE). **A**: We considered populations that have recently diverged with gene flow between them. Some genetic variants will have a different effect on fitness in the diverged population (*s*_2_) than in the ancestral population (*s*_1_). **B**: The joint DFE is defined over pairs of selection coefficients (*s*_1_, *s*_2_). Insets show the joint AFS for pairs of variants that are strongly or weakly deleterious in each population. In each spectrum, the number of segregating variants at a given pair of allele frequencies is exponential with the color depth. **C**: One potential model for the joint DFE is a bivariate lognormal distribution, illustrated here for strong correlation. **D**: We focus on a model in which the joint DFE is a mixture of components corresponding to equality (*ρ* = 1) and independence (*ρ* = 0) of fitness effects. **E**: As illustrated by these simulated allele frequency spectra, stronger correlations of mutation fitness effects lead to more shared polymorphism. Here *w* is the weight of the *ρ* = 1 component in the mixture model.

Little is known about the shape of the joint DFE, so we considered multiple parametric models. The best fit DFEs for single populations tend to be lognormal or gamma distributions (Boyko et al. 2008), although discrete distributions may sometimes fit better (Kousathanas and Keightley 2013; Johri et al. 2020) We first considered a bivariate lognormal distribution (Fig. 1C), because it has an easily interpretable correlation coefficient. However, the bivariate lognormal distribution can be numerically poorly behaved as the correlation coefficient approaches one and the distribution becomes very thin. We also considered another popular probability distribution for modeling DFEs, the gamma distribution, but there are multiple ways of defining a bivariate gamma distribution (Nadarajah and Gupta 2006). We thus focused on a mixture model that consisted of a component corresponding to perfect correlation with weight *w*, and a component corresponding to zero correlation with weight (1 − *w*) (Fig. 1D). To limit the complexity of the model, we assumed that the marginal DFEs were identical for both populations. In this case, the correlation of the overall distribution is equal to the mixture proportion *w*. We thus interpret and discuss *w* as a DFE correlation coefficient.

The DFE correlation profoundly affects the expected AFS (Fig. 1E). Qualitatively, if the correlation is low, there is little shared high-frequency polymorphism. In this case, alleles that are nearly neutral in one population are often deleterious in the other, driving their frequencies lower in that population. If the correlation of the joint DFE is larger, more shared polymorphism is preserved. To calculate the expected AFS for a given demographic model and DFE, we first cached calculations of the expected AFS for a grid of selection coefficient pairs. Assuming independence among sites, the expectation for the full DFE is then an integration over values of *s*_1_, *s*_2_, weighted by the DFE (Fig. S1) (Ragsdale et al. 2016; Kim et al. 2017). We based our approach on the fitdadi framework developed by Kim et al. (2017), and our approach is integrated into our dadi software (Gutenkunst et al. 2009). More detail can be found in the Methods section.

## Simulation

To evaluate the precision of our approach, we first stochastically simulated unlinked single nucleotide polymorphisms (SNPs) under a known demographic model (Table S1 & Fig. S2) and a symmetric lognormal mixture model for the joint DFE (Fig. 1; Eq. 4 & 6). We then inferred the three joint DFE parameters: the mean *µ* and standard deviation *σ* of the marginal lognormal distributions and the DFE correlation *w*. The demographic and joint DFE parameters for these simulations were similar to those we later inferred for human populations under a demographic model of divergence, growth, and migration. When we fit the joint DFE to these simulated data, we found that the variance of the inferred parameters grew only slowly as the sample size decreased (Fig. S3A). This suggests that only modest sample sizes are necessary to confidently infer the joint DFE, similar to how only modest sample sizes are necessary to infer the mean and variance of the univariate DFE (Keightley and Eyre-Walker 2010).

Because our inference approach focuses on shared variation, we expected precision to depend on the divergence time between the populations. To test this, we simulated data sets with sample size similar to our real Drosophila data and varied the divergence time in the demographic model. We found that the variance of the inferred *µ* and *σ* parameters was always small (Fig. S3B & C), but the variance of the inferred DFE correlation *w* depended on the divergence time (Fig. S3B & C). That variance was large for small divergence times (*T* = 10^− 4^). This is expected, because in this case selection has had little time act differently in the two populations. That variance was also large if the divergence time was large and there was no migration between the populations (Fig. S3C). This is also expected, because in this scenario there is little shared variation between populations. However, the variance of the inferred DFE correlation *w* was small when the divergence time was between 10^− 3^ and 10^0^ (Fig. S3B & C).

Having found good precision for our inference, we then turned to testing the robustness of our inference to model misspecification. Since these tests focused on biases in the average inference, we did not stochastically sample data for these analyses, but rather used the expected AFS under each scenario as the data.

The demographic model is a key assumption of our joint DFE inference procedure. To test how imperfect modeling of demographic history would bias our inference, we simulated both neutral and selected data under a demographic model that included divergence, exponential growth in both populations, and asymmetric migration between populations (Fig. S2B). We then fit models that either lacked migration or that modeled instantaneous growth and symmetric migration to the neutral data (Fig. S2C). We then used these mis-specified models to infer the DFE correlation *w* from the selected data. For both misspecified demographic models, although the inferred *µ* and *σ* were biased, we found that the inferred *w* was not strongly biased, particularly for large correlations (Fig. 2A).

**Figure 2:**
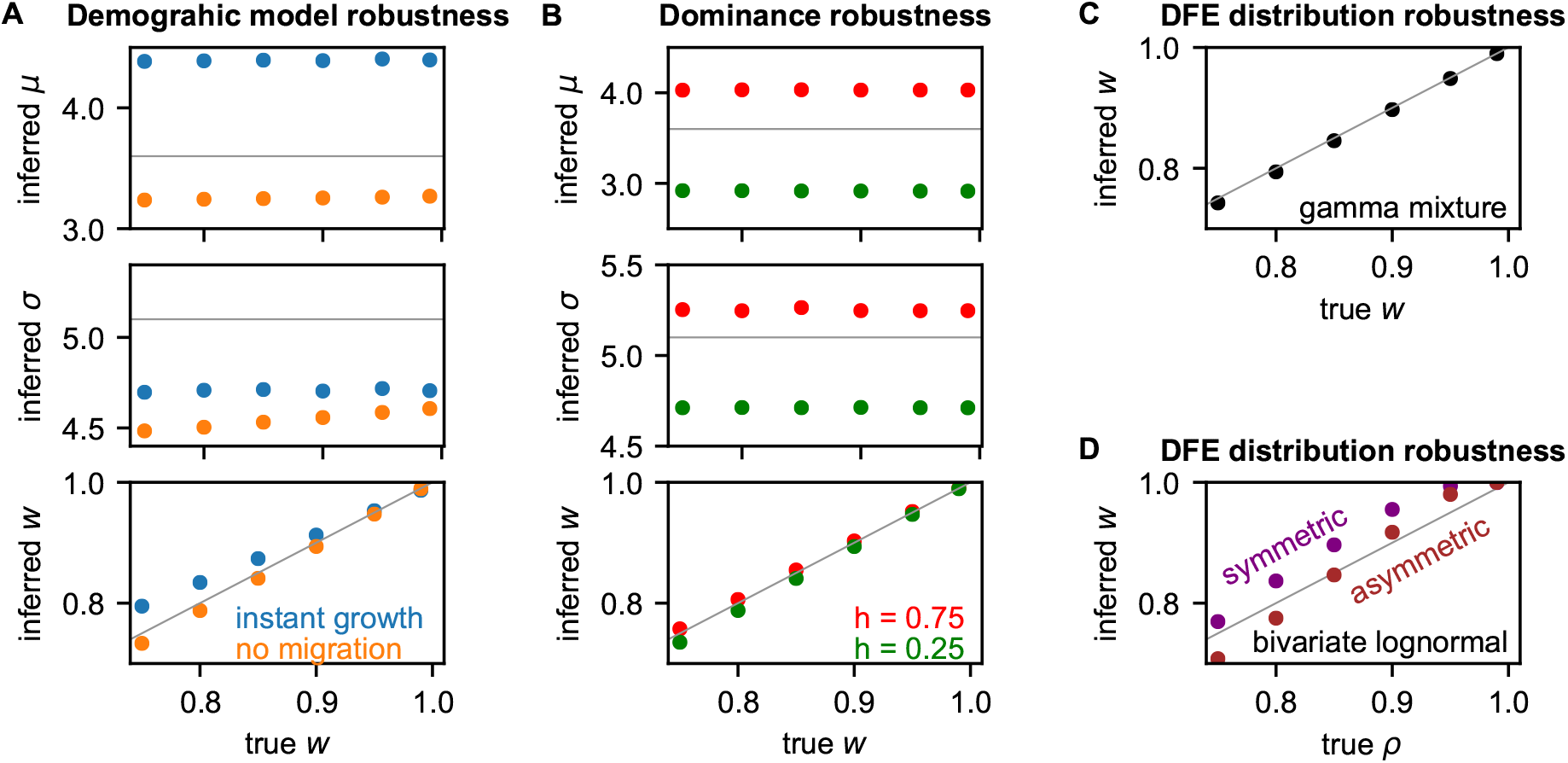
Robustness of joint DFE inference to model misspecification. Simulated neutral and selected data were generated under a demographic model with exponential growth and migration (Table S1), and lognormal mixture DFE models were fit to the data. The DFE parameters are: *µ*, the mean log population-scaled selection coefficient; *σ*, the standard deviation of those log coefficients; and *w*, the correlation of the DFE. The gray lines indicate true values, and the data plotted in these figures can be found in Table S4–S6. **A**: In this case, simpler demographic models with instantaneous growth or symmetric migration were fit to the neutral data. The resulting misspecified model was then used when inferring the DFE. This misspecification strongly biased *µ* and *σ*, but not *w*. **B**: In this case, selected data were simulated assuming dominant or recessive mutations, but the DFE was inferred assuming no dominance (*h* = 0.5). Again, *µ* and *σ* are strongly biased, but *w* is not. **C**: In this case, selected data were simulated using a mixture of gamma distributions. When these data were fit using our mixture of lognormal distributions, *w* was not biased. **D**: In this case, selected data were simulated using bivariate lognormal models, with either symmetric or asymmetric marginal distributions. When these data were fit using our symmetric mixture of lognormal distributions, *w* was only slightly biased.

Dominance is a potential confounding factor when inferring the joint DFE, since dominance influences allele frequencies differently in populations that have and have not undergone a bottleneck (Balick et al. 2015). Typically, mutation fitness effects in diploids are assumed to be additive, corresponding to a dominance coefficient of *h* = 0.5. To test the effects of dominance on our inference, we simulated nonsynonymous frequency spectra with dominance coefficients of *h* = 0.25 and *h* = 0.75 and then optimized joint DFE parameters under the assumption that *h* = 0.5. We found that an incorrect assumption about dominance did not substantially bias the inferred *w*, although it did bias the inferred *µ* and *σ* (Fig 2B).

The probability distribution assumed for the joint DFE is another potential confounding factor. To test how this might bias inference, we first simulated a true mixture model in which the marginal distributions were gamma (Eq. 7), rather than lognormal (Eq. 6). In this case, we found that inferred *w* was not substantially biased (Fig. 2C). We also considered fitting the lognormal mixture model (Fig. 1D) to data simulated under a bivariate lognormal model (Fig. 1C & Eq. 8). In this case, we found that the inferred mixture component *w* was larger than the simulated bivariate lognormal correlation coefficient *ρ*, although they were similar (Fig. 2D). The mixture model assumes symmetric marginal distributions between the two populations, but the bivariate lognormal model is more general and permits asymmetric marginal distributions. When we simulated data under a bivariate model with asymmetric means and variances of the marginal distributions, but fit with a symmetric mixture model, we found only slight bias, similar to the symmetric bivariate case (Fig. 2D).

Finally, background selection may also bias our joint DFE inference. To examine the effects of background selection on our inference, we simulated data with linkage using SLiM 3 (Haller and Messer 2019). We simulated genome-scale data for both human- and Drosophila-like scenarios using the best fit demographic models we inferred for our real data (Fig. S5A & B). For each data set, we fit a demographic model to the simulated synonymous mutations then used that demographic model to infer the joint DFE from the simulated nonsynonyous mutations. For human-like simulations, we also carried out the analysis using simpler demographic models in the inference. As expected, we found that background selection (BGS) biased our demographic model inferences. For example, if we used the same human demographic model in the inference and simulation, the inferred divergence time increased as the DFE correlation *w* decreased (Table S7). As *w* decreased, the strength of BGS increased (Table S8). However, we found that the joint DFE correlation *w* could be robustly inferred in the presence of background selection (Fig. 3). The inferred *µ* and *σ* were biased if the demographic model was misspecified (Fig. 3A). But the inferred *w* was somewhat overestimated only if *w* was less than 0.8 with misspecified demographic models (Fig. 3A). In our Drosophila-like simulations, no bias in inferred *w* was observed (Fig. 3B). Because the strength of background selection in our simulations was much stronger than estimated in empirical studies (Fig. S4), these simulations suggest that our analysis of the real data is robust to background selection.

**Figure 3:**
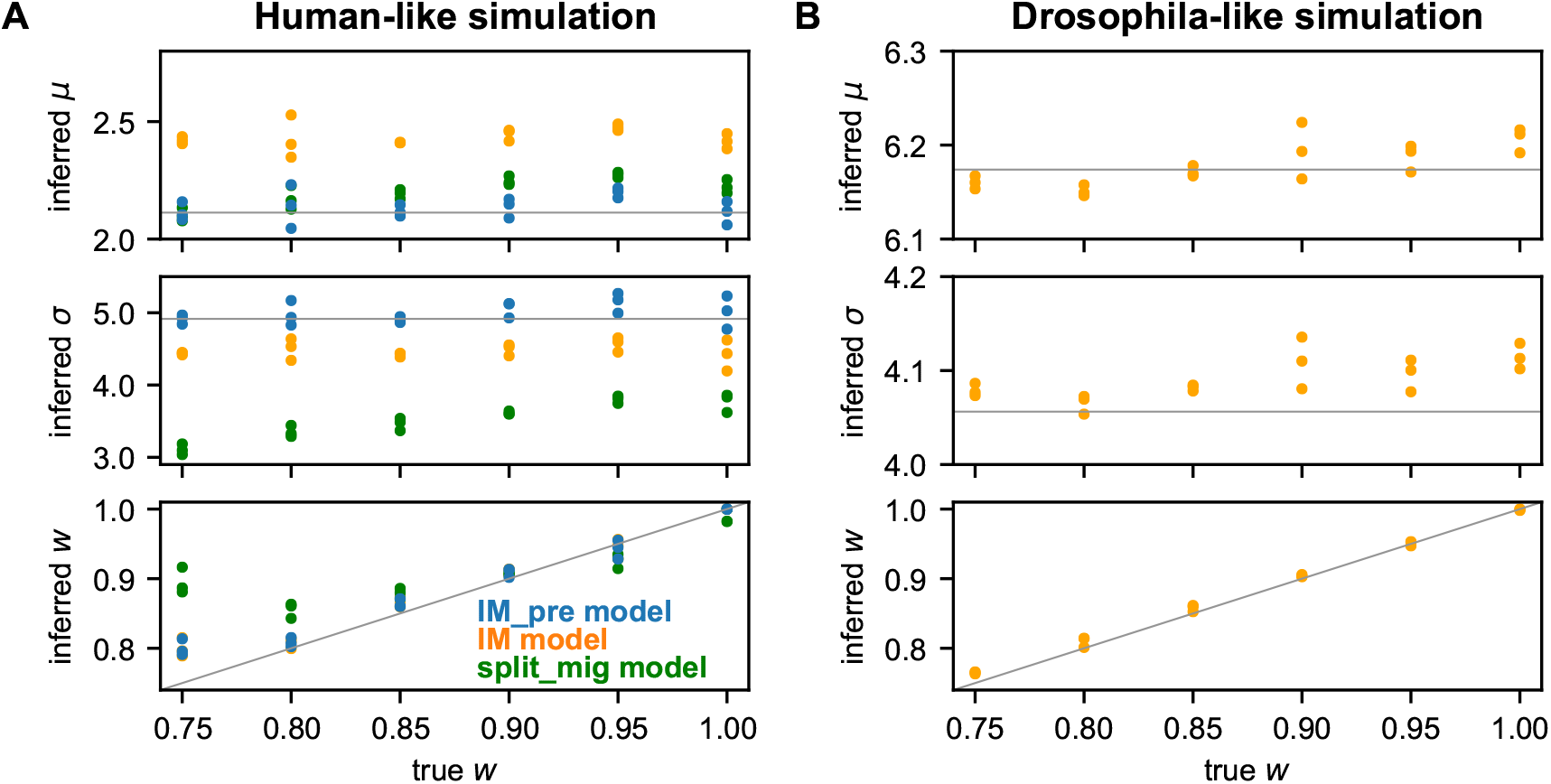
Robustness of joint DFE inference to background selection. Simulated genome-scale data were generated with background selection and different DFE correlations. **A**: Data were simulated using the best fit demographic model for humans in Fig. S5A with *µ* = 2.113 and *σ* = 4.915. Beside fitting the true model, simpler demographic models (Fig. S2) were also fitted to test the robustness of model misspecification in the presence of background selection. **B**: Data were simulated using the best fit demographic model for *D. melanogaster* in Fig. S5B with *µ* = 6.174 and *σ* = 4.056. Points indicate inferences from distinct data sets and colors indicate different demographic models using in the inference. Gray lines indicate true values. The data plotted in these figures can be found in Table S7.

Together, our tests on simulated data suggest that inferring the DFE correlation *w* from the joint AFS can be done with high precision and is robust to multiple confounding factors, including misspecification of the demographic model and DFE distribution as well as the presence of background selection.

### Application

We applied our joint DFE inference approach to humans, *D. melanogaster*, and wild tomatoes. For humans, we considered the joint DFE between Yoruba in Ibadan (YRI) and Utah residents (CEPH) with Northern and Western European ancestry (CEU) populations, because the Yoruba are a well-studied proxy for the ancestral human population and European populations parallel the history of French *D. melanogaster*. For *D. melanogaster*, we considered the joint DFE between Zambian and French populations, because the Zambian population is representative of the ancestral population (Lack et al. 2015) and France is a distinct environment. For wild tomatoes, we considered the joint DFE between two closely related species, *Solanum chilense* and *Solanum peruvianum*, because they still share substantial polymorphism and have overlapping ranges.

We first fit demographic models to synonymous variants in each population pair. For all the three species, we fit relatively simple models of divergence with exponential growth and gene flow, although for humans we also found it necessary to include pre-divergence population growth. Broadly, these models fit the data well (Fig. S5 & S6).

We next estimated the joint DFE using all nonsynonymous variants in the whole exome data from each species with our lognormal mixture model (Fig. 1D). In all the cases, the resulting models fit the nonsynonymous joint frequency spectrum well, with similar patterns of residuals to the demographic models fit to synonymous data (Fig. 4 & S6). For humans, we found the highest DFE correlation *w* = 0.995 *±* 0.007 in our study (Fig. 5 & Table S9), which was statistically indistinguishable from perfect correlation *w* = 1. *±* For *D. melanogaster*, we found that mutation fitness effects between Zambian and French populations were highly correlated, with *w* = 0.967 *±* 0.017 (Fig. 5 & Table S10). For wild tomatoes, we found the lowest DFE correlation, *w* = 0.905 *±* 0.015 (Fig. 5 & Table S11). We also inferred DFE correlations using mutations from non-CpG regions in humans and *D. melanogaster*, because the mutation rate may differ between non-CpG and CpG regions. The resulting estimates of *w* (Fig. S10 & Table S9 & S10) were statistically indistinguishable from those using the whole exome data. Among these three population pairs, the inferred DFE correlation was negatively related to genetic divergence, as measured by *F*_*ST*_ (Fig. 5A).

**Figure 4:**
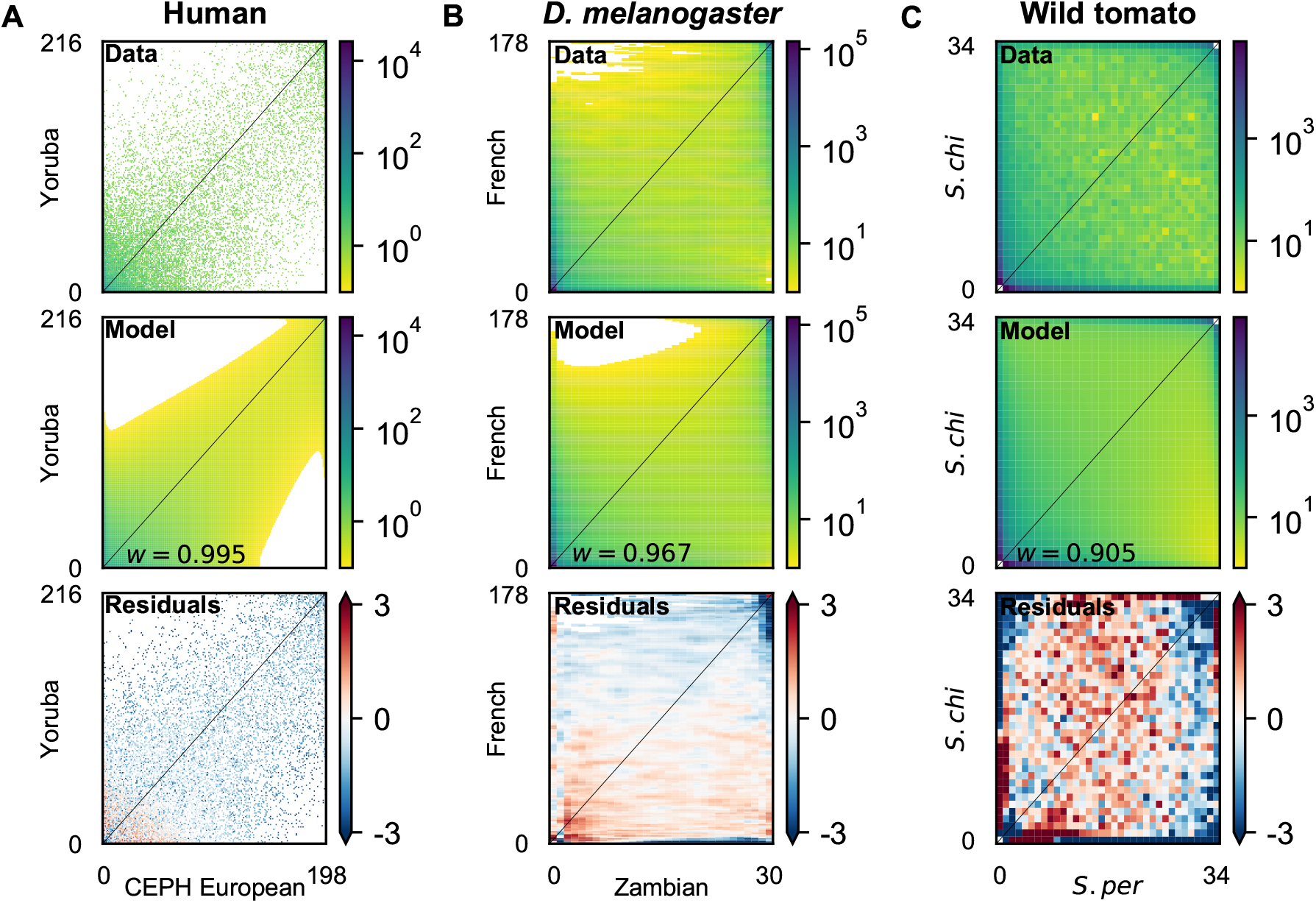
Model fits to joint allele frequency spectra (AFS) using nonsynonymous data. **A**: Joint AFS for the human nonsynonymous data, the best fit model with DFE correlation *w* = 0.995, and the residuals between model and data. **B**: Joint AFS for the *D. melanogaster* nonsynonymous data and the best fit model with DFE correlation *w* = 0.967. **C**: Joint AFS for the wild tomato nonsynonymous data and the best fit model with DFE correlation *w* = 0.905.

**Figure 5:**
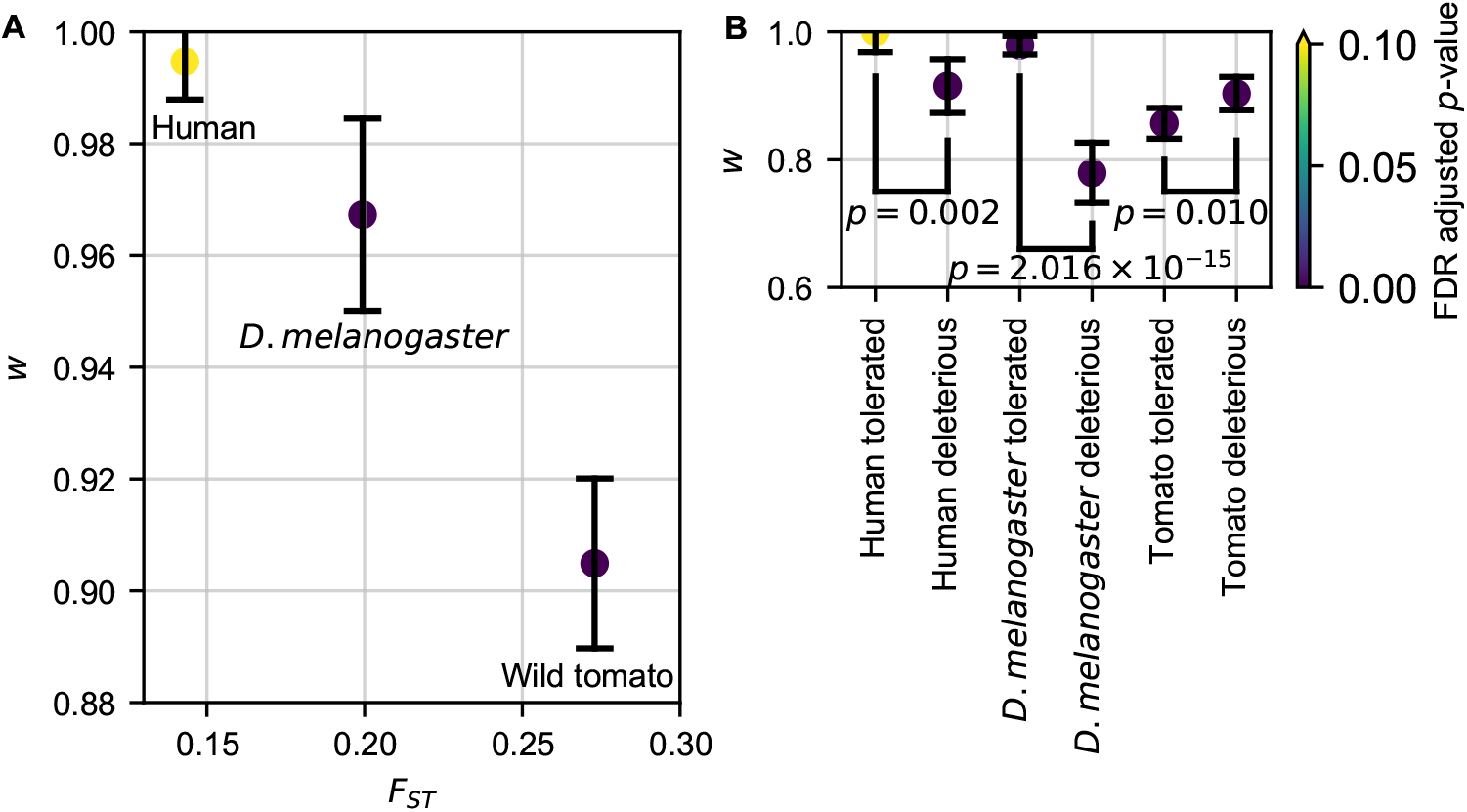
Exome-wide DFE correlations. **A**: Plotted are maximum likelihood inferences of the DFE correlation *w* with 95% confidence intervals versus genetic divergence *F*_*ST*_ of the considered population pair. **B**: Plotted are maximum likelihood inferences of the DFE correlation *w* with 95% confidence intervals for nonsynonymous SNPS with different predicted effects from SIFT. Colors indicate FDR adjusted *p*-values from two-tailed *z*-tests as to whether the confidence interval overlaps *w* = 1. *F*_*ST*_ was estimated using whole-exome synonymous mutations.

For simplicity, we assumed that the DFE correlation *w* is constant throughout the distribution, but the correlation may depend on how deleterious the mutation is. To test this assumption, rather than adding complexity to the DFE model, we instead segregated our data by applying SIFT scores to predict whether a nonsynonymous mutation is likely to be tolerated or deleterious based on evolutionary conservation (Vaser et al. 2016). We then fit DFE models to the SNPs in each class. As expected, we inferred a more negative mean fitness effect for the deleterious class than the tolerated class (Fig. S10 & Table S9–S11). Moreover, we found that the DFE correlation *w* was dramatically smaller for the deleterious class than the value from the tolerated class in humans and *D. melanogaster*, but not in wild tomatoes (Fig. 5B). To test whether this effect extended beyond individual mutations to whole genes, we also separated our data by the dN/dS ratio in humans and *D. melanogaster*. We found no significant difference in DFE correlations among genes with different dN/dS ratios (Fig. S10). However, we did observe that the average strength of purifying selection increases as the dN/dS ratio decreases (Fig. S10).

To investigate the biological basis of the joint DFE, we considered genes of different function based on Gene Ontology (GO) terms (The Gene Ontology Consortium 2000). For *D. melanogaster*, we found a wide range of inferred DFE correlations, with the lowest maximum likelihood estimate corresponding to mutations in genes involved in the mitotic nuclear division at *w* = 0.901 *±* 0.048 (Fig. 6 & Table S10). For wild tomatoes, we found an even wider range of inferred DFE correlations, with the lowest maximum likelihood estimate being genes involved in photosynthesis at *w* = 0.769 *±* 0.106 (Fig. 6 & Table S11). For humans, we found that all GO terms yielded values of *w* that were statistically indistinguishable from one (Table S9 & Fig. S7). Among the *D. melanogaster* GO terms, we found no correlation between the inferred *w* and the mean and standard deviation of the marginal DFEs (Fig. S11), suggesting that the variation we see in *w* is not driven simply by variation in overall constraint. In humans, we further explored the biological context of the joint DFE by considering genes that are involved in disease and that interact with viral pathogens. We found no statistically significant differences in DFE correlations among these gene groups, although we did find that the DFE for genes involved in disease or that interact with viruses was shifted toward more negative selection (Table S9 & Fig. S10).

**Figure 6:**
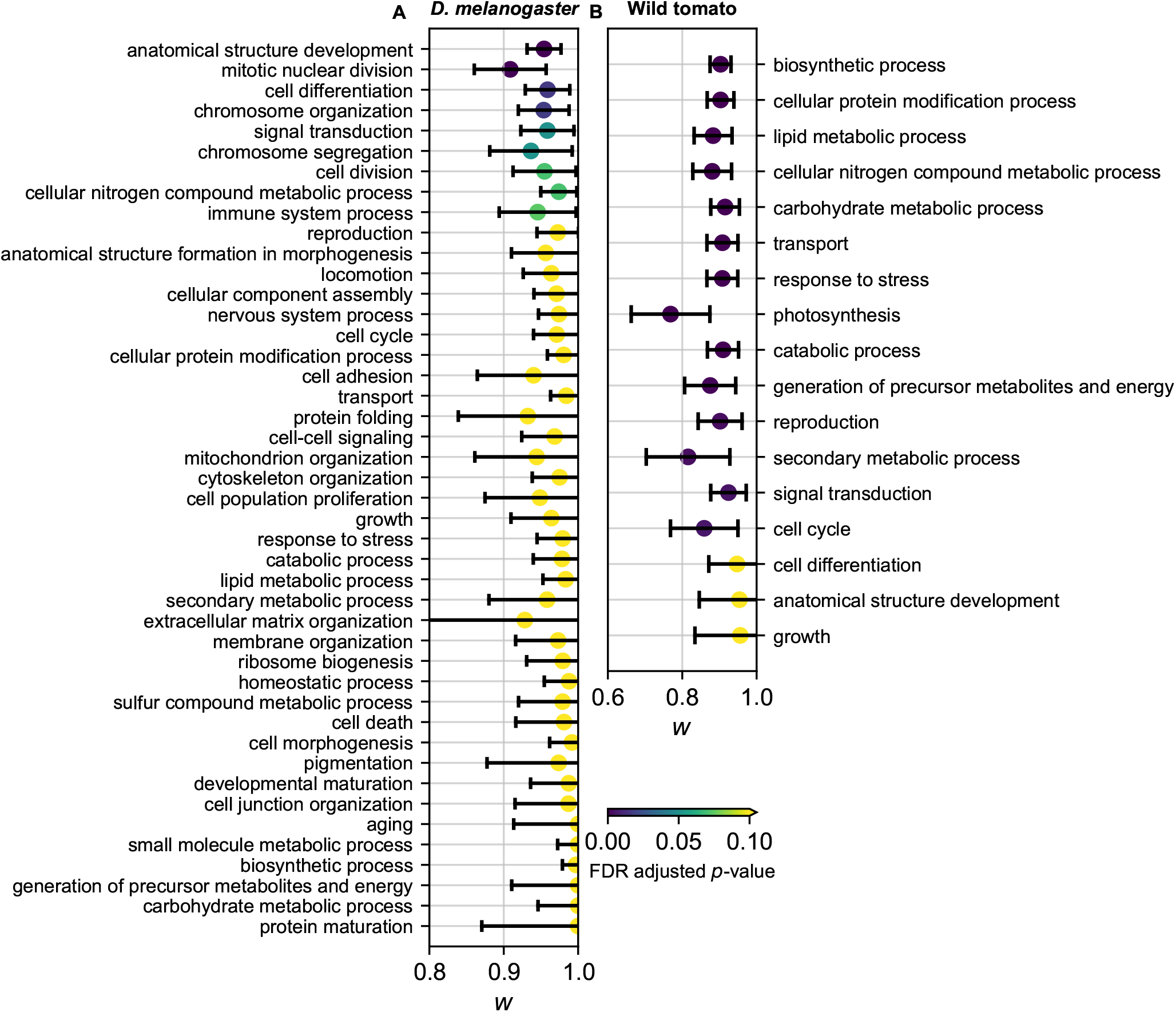
DFE correlation for different GO terms in *D. melanogaster* and wild tomatoes. Plotted are maximum likelihood inferences with 95% confidence intervals. Colors indicate FDR-adjusted *p*-values from two-tailed *z*-tests as to whether the confidence interval overlaps *w* = 1. The data plotted in these figures can be found in Table S10 & S11. **A**: Inferred DFE correlation in *D. melanogaster*. **B**: Inferred DFE correlation in wild tomatoes.

To test the robustness of our analyses in the real data to various modeling choices, we used the variation among our inferences among *D. melanogaster* GO terms. We fit simpler models of demographic history with instantaneous growth in the two diverged populations with and without symmetric migration to the synonymous data and used those models as the basis of joint DFE analysis. Although these demographic models fit the data much less well than our main model (Fig. S6 & S12), the inferred values of *w* for the GO terms were highly correlated with those from our main model (Fig. S13A & B). We also tested our approach using a DFE model with a bivariate lognormal model instead of a lognormal mixture model. The inferred values for *ρ* in the bivariate model were highly correlated with the values for the inferred *w* (Fig. S13C). Together, these results suggest that the robustness we observed in simulated data (Fig. 2) holds true for real data.

## Discussion

In this study, we introduced the concept of a joint distribution of fitness effects between pairs of populations, and we developed and applied an approach for inferring it. We tested our approach with simulation studies and found that inferring the DFE correlation between populations does not require excessive data and is robust to many forms of model misspecification (Fig. S3, 2, & 3). We then applied our approach to humans, *D. melanogaster*, and wild tomatoes. Among these species, we found the lowest exome-wide DFE correlation in wild tomatoes and the highest in humans (Fig. 5A). In humans and *D. melanogaster*, we found that the DFE correlation is lower for deleterious mutations than tolerated mutations (Fig. 5B). And in *D. melanogaster* and tomatoes, we found that the DFE correlation varied with gene function (Fig. 6). These results illustrate the biological insights that can be gained by considering the joint DFE between populations.

The first step of our analyses is fitting a demographic model, although our correlation inferences are robust to details of that model (Fig. 2A & S12). Nevertheless, our inferred demographic models (Fig. S5) agree well with other inferences. For humans, our demographic parameters were similar to those of Gravel et al. (2011). For *D. melanogaster*, our inferred population sizes and divergence time for African and European populations agree with those of Arguello et al. (2019), although we used different populations and different models. For wild tomatoes, we obtained a demographic model close to the result of Beddows et al. (2017).

The fitness effect of a mutation may differ between populations due to differences in both environmental and genetic context. The wild tomato species we analyzed overlap in range and are more genetically differentiated than the *D. melanogaster* or human populations we studied. In this case, we speculate that differences in fitness effects are primarily driven by differences in genetic background, although *Solanum chilense* does exhibit adaptations for more arid habitats (Moyle 2008). Among the species we studied, humans exhibited the highest correlation of mutation fitness effects, which was statistically indistinguishable from perfect correlation *w* = 1, suggesting little difference in mutation fitness effects between YRI and CEU populations. Huang et al. (2021) also estimated the genome-wide differences of selection coefficients between Africans and Europeans were almost 0 with a different approach (He et al. 2015). It is unclear whether this is caused by our relatively low genetic differentiation or our ability to control our local environment. Experiments suggest that stressful environments can alter DFEs between populations (Wang et al. 2014). Previous population genetic studies also have found evidence for differences in marginal DFEs between populations of humans (Boyko et al. 2008; Lopez et al. 2018) and also between populations of other primates (Ma et al. 2013; Castellano et al. 2019; Tataru and Bataillon 2019). Although we assumed that the mean and the variance of mutation fitness effects did not differ between the two populations in our models for the joint DFE, those previous studies found only slight differences and our simulation study suggests that inferences of the DFE correlation are robust to relatively large differences in marginal DFEs (Fig. 2D). Recently, Martin and Lenormand (2015) extended Fisher’s Geometrical Model to consider the relationship between mutation fitness effects in two different environments, represented by two optima in trait space. Unfortunately, they could not derive an analytic joint DFE for their model, so we could not apply it here. Overall, our simple models of the joint DFE fit the data well, but more complex models may be more informative. Over the long term, assessing the joint DFE between multiple populations of multiple species may reveal the relative importance of environmental and genetic context in determining the mutation fitness effects.

We focused on the deleterious component of the DFE in this study, and positive selection or local adaptation may affect joint DFE inference. However, Castellano et al. (2019) found that including beneficial mutations or not did not affect the DFE model for the deleterious components in humans. Moreover, Zhen et al. (2021) estimated the proportion of new beneficial mutations to be around 1.5% in humans and close to 0 in *D. melanogaster*. Therefore, we do not expect beneficial mutations to significantly affect our inference in humans and *D. melanogaster*. Further studies that include local adaptation when inferring the joint DFE may improve our analysis of populations with low DFE correlations, such as wild tomatoes.

Finally, the concept of a joint DFE could be widely applicable. For example, we recently inferred a joint DFE between mutations at the same protein site, using triallelic variants (Ragsdale et al. 2016). Remarkably, we found that biochemical experiments in a variety of organisms yielded a similar correlation of pairwise fitness effects to the value we inferred from *D. melanogaster* population genetic data. Other potential applications of a joint DFE include modeling ancient DNA data to infer DFE correlations across time and modeling linkage to infer DFE correlations across genomic positions. We thus anticipate that extending the concept of the DFE from one population to two or more will significantly advance our understanding of population evolution and have broad impact in population genetics.

## Methods

### Inferring joint DFE from joint allele frequency spectrum (AFS)

If we sample *n*_1_ chromosomes from population 1 and *n*_2_ chromosomes from population 2, then the joint AFS for these two populations can be written as

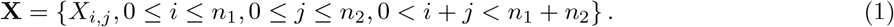

Here, *X*_*i,j*_ denotes the number of mutations in the sample that have *i* copies of derived alleles among the *n*_1_ chromosomes from population 1 and *j* copies of derived alleles among the *n*_2_ chromosomes from population 2. We denote the joint spectra for neutral and selected mutations as **N** = {*N*_*i,j*_} and **S** = {*S*_*i,j*_}, respectively.

Let **F**(*γ*_1_, *γ*_2_|Θ_demo_) = {*F*_*i,j*_(*γ*_1_, *γ*_2_|Θ_demo_)} be the expected joint AFS for demographic parameters Θ_demo_, population-scaled selection coefficients *γ*_1_ in the ancestral and first contemporary population and *γ*_2_ in the second contemporary population, and population-scaled mutation rate *θ* = 1. Then the expected neutral joint AFS is

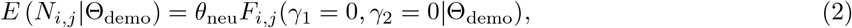

where *θ*_neu_ is the population-scaled neutral mutation rate (Gutenkunst et al. 2009). The expected selected joint AFS is

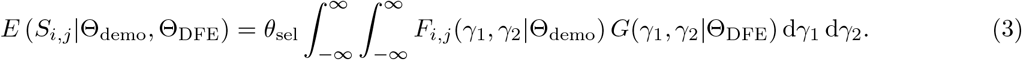

Here *θ*_sel_ is the population-scaled mutation rate for selected mutations, and *G*(*γ*_1_, *γ*_2_|Θ_DFE_) is the joint DFE.

In most of our analyses, we modeled the joint DFE as a mixture of two components, *G*_1d_ and *G*_2d_, where *G*_1d_ is a DFE with equal selection coefficients in the two populations, and *G*_2d_ is a DFE with statistically independent selection coefficients and marginal distributions *G*_1d_. Letting *w* be the mixture proportion of*G*_1d_, we have

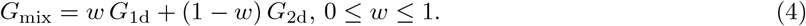

And considering only deleterious mutations we have

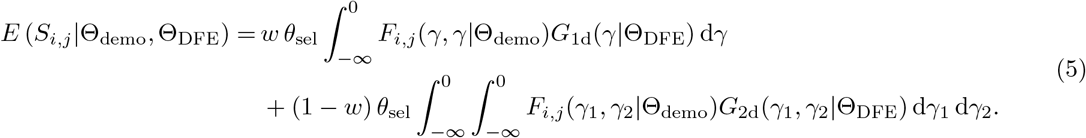

We typically worked with lognormal distributions, so

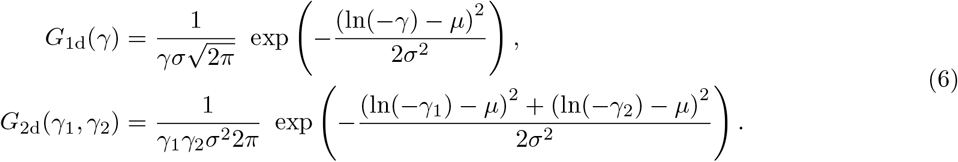

Here *µ* and *σ* are the mean and standard deviation of the logs of the population-scaled selection coefficients, respectively.

To test the robustness of our approach, we also considered other models for the joint DFE. When using a mixture of gamma distributions,

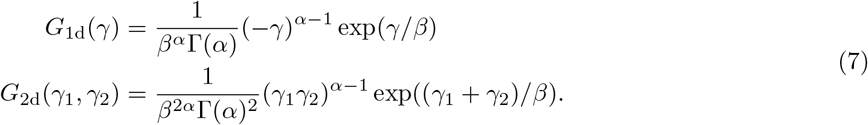

Here *α* is the shape parameter and *β* is the scale parameter. When using a bivariate lognormal distribution, which is potentially asymmetric,

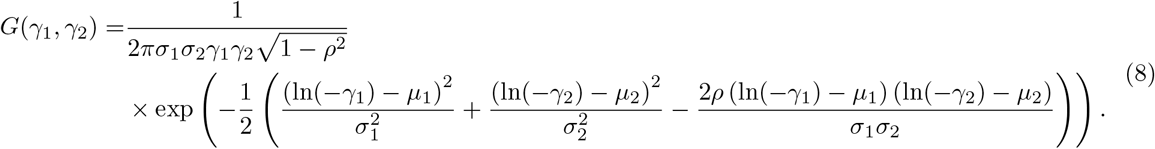

Here *ρ* is the correlation coefficient.

Calculating the expected selected joint AFS (Eq. 3 & 5) is computationally expensive, because spectra **F**(*γ*_1_, *γ*_2_|Θ_demo_) must be calculated for many pairs of selection coefficients. Simultaneously inferring the demographic Θ_demo_ and the DFE Θ_DFE_ parameters is thus infeasible. We thus first inferred the demographic parameters using the putative neutral data and then held those parameters constant while inferring the DFE parameters.

We inferred the demographic parameters 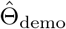 by maximizing the composite likelihood of the neutral joint AFS, including *θ*_neu_ as a free parameter (Gutenkunst et al. 2009). To then infer the DFE parameters Θ_DFE_, we modeled the selected joint AFS as a Poisson Random Field (Sawyer and Hartl 1992) and maximized the composite likelihood

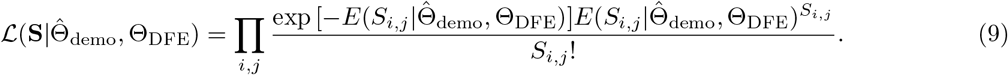

Here, 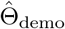 represents the demographic parameters inferred from the neutral data. And in this step we fixed *θ*_sel_ to a multiple of *θ*_neu_ determined by the expected ratio of new selected to new neutral mutations, based on base-specific mutation rates and genome composition.

Numerically, to calculate the expected selected joint AFS, we first cached expected spectra 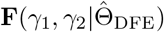 for a range of selection coefficient pairs. The cached values of *γ*_1_, *γ*_2_ were from 50 points logarithmically spaced within [− 10^− 4^, − 2000], for a total of 2500 cached spectra (Fig. S1). We then evaluated Eq. 3 using the trapezoid rule over these cached points. For the mixture model (Eq. 5), the *G*_1d_ component was calculated as a one-dimensional integral over a cache of *γ*_1_ = *γ*_2_ spectra. Probability density for the joint DFE may extend outside the range of cached spectra. To account for this density, we integrated outward from the sampled domain to *γ* = 0 or − ∞ to estimate the excluded weight of the joint DFE. We then weighted the closest cached joint AFS **F** by the result and added it to the expected joint AFS. For the edges of the domain, this was done using the SciPy method quad, and for the corners it was done using dblquad (Virtanen et al. 2020).

### Simulated data

For our precision tests (Fig. S3), we used dadi to simulate data sets without linkage. Unless otherwise specified, for Fig. S3 and Fig. 2, the “truth” simulations were performed with an isolation-with-migration (IM) demographic model (Fig. S2B) with parameters as in Table S1, a joint lognormal mixture DFE model with marginal mean *µ* = 3.6 and standard deviation *σ* = 5.1, and with sample sizes of 216 for population 1 and 198 for population 2. For Fig. S3, data were simulated with *w* = 0.9 and the nonsynonymous population-scaled mutation rate *θ*_*NS*_ = 13842.5 by Poisson sampling from the expected joint AFS. For Fig. S3A the resulting average number of segregating polymorphisms varied with sample size, ranging from 6,953 for sampling two chromosomes to 45,691 for sampling 100 chromosomes. For Fig. S3B & C, the sample size was fixed at 20 chromosomes per population.

For our robustness tests (Fig. 2), we were interested in bias rather than variance, so misspecified models were fit directly to the expected frequency spectrum under the true model without Poisson sampling noise. For Fig. 2A, the best fit model with no migration had *s* = 0.937, _1_ = 3.025, _2_ = 3.219, *T* = 0.0639, *m* = 0, and the best fit model with instantaneous growth and symmetric migration had _1_ = 2.4, _2_ = 0.92, *T* = 0.23, *m* = 0.42. For Fig. 2C, the true joint DFE was a mixture model with marginal gamma distributions with *α* = 0.4, *β* = 1400. For Fig. 2D, the true joint DFE was a symmetric bivariate lognormal distribution with *µ* = 3.6 and *σ* = 5.1, and for the asymmetric case in Fig. 2D, *µ*_1_ = 3.6, *σ*_1_ = 5.1, *µ*_2_ = 4.5, *σ*_2_ = 6.8. We then simulated data with different correlation coefficients *ρ* to examine the relationship between *ρ* and the DFE correlation *w*.

To examine the effects of background selection, we used SLiM 3 (Haller and Messer 2019) to simulate data with linkage. We replicated our simulation and inference three times for each *w* with different demographic models in the human simulations and an IM model in the *D. melanogaster* simulations (Fig. S2). For humans, we simulated the exome in chromosome 21 using the demographic parameters in Fig. S5A, the joint

DFE parameters *µ* and *σ* from the whole human exome in Table S9 with *w* = 0.75, 0.8, 0.85, 0.9, 0.95, 1, and sample sizes of 216 for population 1 and 198 for population 2. We assumed the mutation rate was 1.5 × 10^− 8^ per nucleotide per generation (Ségurel et al. 2014). We further assumed the ratio of the nonsynonymous to synonymous mutations in humans was 2.31 (Huber et al. 2017). In our simulation, we used the human exome based on the reference genome hg19 from UCSC Genome Browser and the deCODE human genetic map (Kong et al. 2010). For each *w*, we first simulated human chromosome 21 twenty times, then obtained 20 synonymous frequency spectra and 20 nonsynonymous frequency spectra from these sequences. We combined these 20 synonymous frequency spectra into a single one and inferred the demographic models. We then combined the 20 nonsynoymous frequency spectra into one spectrum and inferred the joint DFEs. We inferred the joint DFEs using both the true (IM pre model) and wrong (IM model with asymmetric migration & split mig model without migration) demographic models (Fig. S2). For *D. melanogaster*, we simulated small sequences instead of a whole chromosome, because the large population size of *D. melanogaster* made our simulation extremely slow. We used the demographic parameters for the IM model in Fig. S5B, the joint DFE parameters *µ* and *σ* from the whole *D. melanogaster* exome in Table S10 with *w* = 0.75, 0.8, 0.85, 0.9, 0.95, 1, and sample sizes of 178 for population 1 and 30 for population 2. For each *w*, we simulated 20,000 small sequences with a length of 1000 bp, then obtained 20,000 synonymous frequency spectra and 20,000 non-synonymous frequency spectra. We combined these 20,000 synonymous frequency spectra into a single one and inferred the demographic models. We then combined the 20,000 nonsynonymous frequency spectra into one spectrum and inferred the joint DFEs. This was equivalent to a total sequence size of 20 Mb. We assumed that the mutation rate was 2.8 × 10^− 9^ per nucleotide per generation and that the recombination rate was 5 × 10^− 9^ per nucleotide per generation (Keightley et al. 2014). We also assumed the ratio of the nonsynonymous to synonymous mutations in *D. melanogaster* was 2.85 (Huber et al. 2017). To accelerate our simulation, we used a factor of 1000 to re-scale the population size, mutation rate, and recombination rate (Hoggart et al. 2007). To quantify the strength of BGS in our simulations, we simulated data under neutral models and compared the expected number of pairwise differences between two chromosomes in the non-neutral scenarios with the neutral ones (Hudson and Kaplan 1995). The strength of BGS (Fig. S4) in the simulated data for both humans and *D. melanogaster* was stronger than the estimated strength from the empirical studies (Charlesworth 2013).

### Genomic data

In all analyses, we only considered biallelic SNPs from automosomes. For humans, we obtained 108 and 99 unrelated individuals (216 and 198 haplotypes) from YRI and CEU populations in the 1000 Genomes Project Phase 3 genotype data (The 1000 Genomes Project Consortium 2015). We removed those regions that were not in the 1000 Genomes Project phase 3 strict mask file. We only considered biallelic exonic SNPs that were annotated as synonymous variant or missense variant by the 1000 Genomes Project. We further excluded SNPs without reported ancestral alleles. We also used the CpG table from the UCSC Genome Browser to distinguish SNPs in CpG regions.

For *D. melanogaster*, we obtained Zambian and French *D. melanogaster* genomic data from the Drosophila Genome Nexus (Lack et al. 2016). The Zambian sequences were 197 haploids from the DPGP3 and the French were 87 inbred individuals. We removed those SNPs in the IBD and/or admixture masks. In these data, many SNPs were not called in all individuals. We thus projected downward to obtain a consensus AFS with maximal genome coverage. For these data, that was to a sample size of 178 Zambian and 30 French haplotypes (Fig.S14). We used *D. simulans* as the outgroup and downloaded the alignment between the reference genome for *D. simulans* (drosim1) and the reference genome for *D. melanogaster* (dm3) from UCSC Genome Browser to determine the ancestral allele of each SNP. We then used GATK (version: 4.1.4.1) (McKenna et al. 2010) to liftover the genomic coordinates from dm3 to dm6 with the liftover chain file from the UCSC Genome Browser. To annotate SNPs to their corresponding genes and as synonymous or nonsynonymous mutations, we used ANNOVAR (version: 20191024) (Wang et al. 2010) with default settings and the dm6 genome build. We downloaded the CpG table from the UCSC Genome Browser to distinguish SNPs in CpG regions.

For wild tomatoes, we obtained *Solanum chilense* and *Solanum peruvianum* DNA sequencing data from Beddows et al. (2017) and followed their scheme for assigning individuals to species. We only analyzed 17 *Solanum chilense* and 17 *Solanum peruvianum* individuals sequenced by Beddows et al. (2017) because of their high quality. We used an *Solanum lycopersicoides* individual sequenced by Beddows et al. (2017) to determine the ancestral allele of each SNP. We further removed variants with heterozygous genotype in this *Solanum lycopersicoides* individual. To more easily apply SIFT, we used the NCBI genome remapping service to convert the data from SL2.50 coordinates to SL2.40.

### Fitting demographic models to genomic data

We used dadi to fit models for demography to spectra for synonymous mutations (Gutenkunst et al. 2009), including a parameter for ancestral state misidentification (Ragsdale et al. 2016). For the human analysis, we used dadi with grid points of [226,236,246], and we found that an isolation-with-migration model with an instantaneous growth in the ancestral population (IM pre) fit the data well (Fig. S5A). For the *D. melanogaster* analysis, we used dadi with grid points of [188,198,208], and we found that an IM model fit the data well (Fig. S5B). For the wild tomato analysis, we used dadi with grid points of [44,54,64] and fit a split-migration model with asymmetric migration (Fig. S5C), as Beddows et al. (2017) did.

### Fitting joint DFEs to genomic data

Cached allele frequency spectra were created for the corresponding demographic models. For humans and *D. melanogaster*, we used the same grid points settings as the grid points used when inferring demographic models. For wild tomatoes, we used dadi with grid points of [300, 400, 500] to generate caches with selection. Models of the joint DFE were then fit to nonsynonymous data by maximizing the likelihood of the data, assuming a Poisson Random Field (Sawyer and Hartl 1992). In these fits, the population-scaled mutation rate for nonsynonymous mutations *θ*_*NS*_ was held fixed at a given ratio to the rate for synonymous mutations *θ*_*S*_ in the same subset of genes, as inferred from our demographic history model. For *D. melanogaster* this ratio was 2.85 and for humans it was 2.31 (Huber et al. 2017). For wild tomatoes, this ratio was assumed to be 2.5, which was between the ratios in humans and *D. melanogaster*. For the lognormal mixture model, the three parameters of interest are the DFE correlation *w* as well as the mean *µ* and standard deviation *σ* of the marginal distributions. In addition, we included a separate parameter for ancestral state misidentification for each subset of the data tested, because the rate of misidentification depends on the strength of selection acting on the sites of interest. To mitigate the effect of background selection, we separately inferred demographic parameters for each subset of the data (Table S9–S11) with the best fit demographic model inferred from the whole exome data (Fig. S5).

We separately analyzed SNPs from genes associated with different Gene Ontology (GO) terms. We downloaded the Generic GO subset from http://geneontology.org/docs/download-ontology/ on August 12, 2020. This is a set of curated terms that are applicable to a range of species (The Gene Ontology Consortium 2000). We considered the direct children of GO:0008510 “Biological Process”, and any gene annotated with a child of a given term was assumed to also be annotated by the parent term. Thus, a given gene may be present in multiple GO terms in our analysis. We used Ensembl Biomart (Cunningham et al. 2019) to retrieve the annotated GO terms for each gene. For humans, we downloaded the GO annotation from https://grch37.ensembl.org/biomart/martview/ with Ensembl Genes 101 database and Human genes (GRCh37.p13) on August 19, 2020. For *D. melanogaster*, we downloaded the GO annotation from https://www.ensembl.org/biomart/martview/ with Ensembl Genes 101 database and *D. melanogaster* genes (BDGP6.28) on September 10, 2020. For tomatoes, we downloaded the GO annotation from https://jul2018-plants.ensembl.org/biomart/martview/ with Ensembl Plants Genes 40 database and Solanum lycopersicum genes (SL2.50) on September 26, 2020. To ensure convergence in our inference, we removed those GO terms with less than 2,000 either synonymous or nonsynonymous mutations (Table S9–S10).

We also separately analyzed SNPs classified by SIFT as deleterious (SIFT score ≤ 0.05) or tolerated (SIFT score > 0.05) (Vaser et al. 2016). We downloaded SIFT predictions from https://sift.bii.a-star.edu.sg/sift4g/ on October 2, 2020. We used the SIFT prediction data with GRCH37.74 for humans, with BDGP6.83 for *D. melanogaster*, and with SL2.40.26 for tomatoes. To carry out our DFE analysis, we needed to estimate an appropriate population-scaled nonsynonymous mutation rate *θ*_*NS*_ for deleterious and tolerated mutations. To do so, we estimated the proportions of deleterious and tolerated mutations in the downloaded SIFT prediction datasets. This is because all the possible mutations and their SIFT scores were predicted in the downloaded datasets. We then obtained the population-scaled mutation rates for deleterious and tolerated mutations by multiplying *θ*_*NS*_ from the whole exome data with the proportions of deleterious and tolerated mutations respectively.

We further considered differences between regions of the genome that experience different levels of evolutionary conservation, as estimated from the ratio of nonsynonymous to synonymous divergence dN/dS. For humans, we separated SNPs into categories based on the estimated dN/dS values of the gene in which they are found from a previous study (Gayà-Vidal and Albà 2014). For *D. melanogaster*, we separated SNPs based on the dN/dS estimate of the surrounding 10 kb genomic region from PopFly (Hervas et al. 2017).

For humans, we also divided genes into classes based on their role in disease and interactions with viruses. Following Struck et al. (2018), we classified genes as associated with Mendelian disease, complex disease, or no disease using Online Mendelian Inheritance in Man (OMIM, Amberger et al. 2015) and the European Bioinformatics Institute’s genome-wide association studies (GWAS) catalog (MacArthur et al. 2017). We used the data of Enard and Petrov (2018) to annotate 4,534 genes as encoding virus-interacting proteins (VIPs). We defined the set of non-VIP genes as the 17,603 Ensembl genes that were not annotated as encoding VIPs. We identified 1,728 genes as known to interact with 2 or more viruses, leaving 2,806 genes known to interact with only a single virus.

To estimate the uncertainty of our inferences, we used an approach based on the Godambe Information Matrix (Coffman et al. 2016), which is computationally more efficient than conventional bootstrap parameter optimization. To generate the requisite bootstrap data sets, we divided the reference genomes into 1 Mb chunks. Because gene content varied among bootstraps, *θ*_*NS*_ also needed to vary. To estimate the appropriate *θ*_*NS*_ for each bootstrap, we scaled corresponding *θ*_*NS*_ from the real data by the ratio of the number of segregating sites in the AFS of the bootstrap versus real data. We found good agreement between the uncertainties estimated by the Godambe approach and those from directly fitting the bootstrap data sets (Fig. S15). Note that this process does not propagate uncertainty in the demographic parameter inference, so our uncertainties are somewhat underestimated.

To estimate *p*-values for inferred DFE correlation *w*, we used the two-tailed *z*-test by assuming *w* = 1 under the null hypothesis and using the standard deviation estimated from the Godambe approach. To compare inferred DFE correlations between tolerated and deleterious mutations, we used two-tailed *z*-tests to calculate *p*-values by assuming no difference under the null hypothesis and using the standard deviations estimated from the Godambe approach. For multiple testing correction, we estimated false discovery rate (FDR) adjusted *p*-values by the Benjamini–Hochberg procedure (Benjamini and Hochberg 1995). These multiple hypothesis tests are from different types of data, including whole-exome data, whole-exome data without CpG regions, different GO terms, genes with different dN/dS values, genes with different SIFT scores, genes associated with no/simple/complex diseases, and genes associated with no/single/multiple VIPs (Table S9–S11).

## Data availability

The software tools used in this study are dadi (https://bitbucket.org/gutenkunstlab/dadi) and SLiM 3 (https://github.com/MesserLab/SLiM/). The supplementary tables can be found at https://doi.org/10.6084/m9.figshare.13936985.

## Acknowledgments

We thank Laura Rose and Thorsten Kloesges for sharing their tomato data and assistance with the analysis. We thank David Enard for sharing his virus-interacting-proteins data. We thank Christian Huber and Kirk Lohmuller for discussing SLiM simulation with background selection. Research reported in this publication was supported by the National Institute of General Medical Sciences of the National Institutes of Health under award number R01GM127348.

## Author contribution

R.N.G designed the study. X.H, A.L.F, A.J.C, T.J.S., M.N.I, J.E.J, J.E.L, A.P.R, and R.N.G designed the analysis method and simulated and analyzed the data. X.H., A.L.F., T.J.S., J.E.J, and R.N.G wrote the manuscript.

## Supporting Information

**Table S1:**
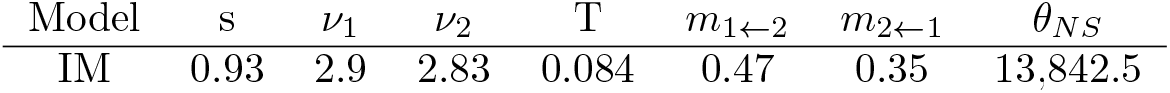
Model parameters for unlinked simulation

**Figure S1:**
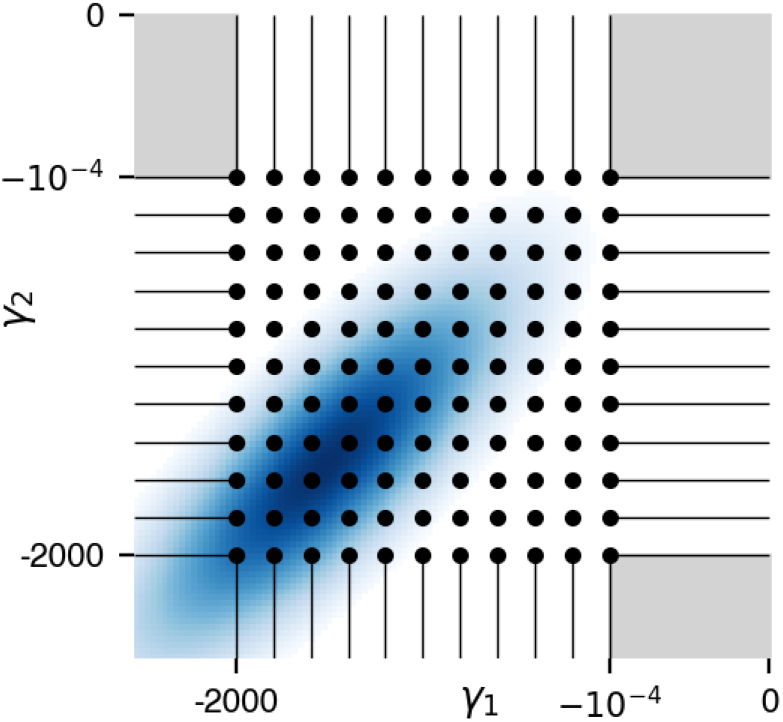
Illustration of computational approach for calculating expected joint AFS for a given joint DFE. Dots represent cached frequency spectra. Horizontal and vertical lines indicate single-variable semi-analytic integrations to estimate DFE density outside the sampled domain, and gray regions indicate corresponding double-variable integrations.

**Figure S2:**
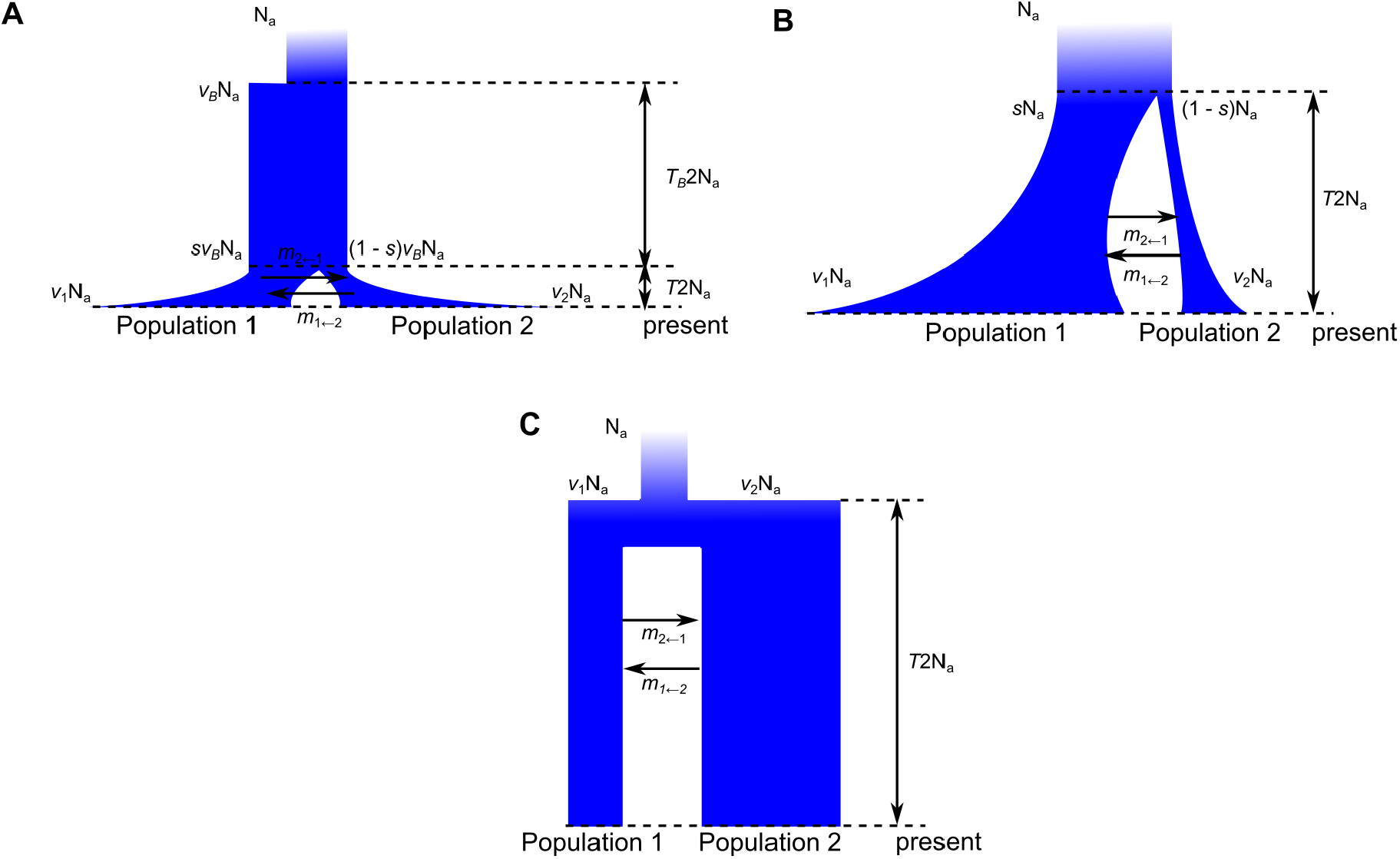
Demographic models used in this study. In these models, the population sizes were scaled by the ancestral population size *N*_*a*_ and the times were in units of 2*N*_*a*_ generations. The migration rates between Population 1 and 2 can be symmetric (*m*_1←2_ = *m*_2←1_) or asymmetric (*m*_1←2_ = *m*_2←1_). If there was no migration, then *m*_1←2_ = *m*_2←1_ = 0. **A**: IM pre model. In this model, the ancestral population experienced an instantaneous population size change at time *T* + *T*_*B*_ before present. After the change, its population size became *ν*_*B*_*N*_*a*_ and remained constant until time *T* before present. Population 1 and 2 diverged at time *T* before present and then grew exponentially. For Population 1, its initial population size was *sν*_*B*_*N*_*a*_ and its final population size was *ν*_1_*N*_*a*_ at present. For Population 2, its initial population size was (1 − *s*)*ν*_*B*_*N*_*a*_ and its final population size was *ν*_2_*N*_*a*_ at present. **B**: IM model. This model corresponds to the IM pre model with no ancestral growth event. **C**: split mig model. In this model, Population 1 and 2 diverged at time *T* before present. Their population sizes remained constant after the divergence with *ν*_1_*N*_*a*_ for Population 1 and *ν*_2_*N*_*a*_ for Population 2.

**Figure S3:**
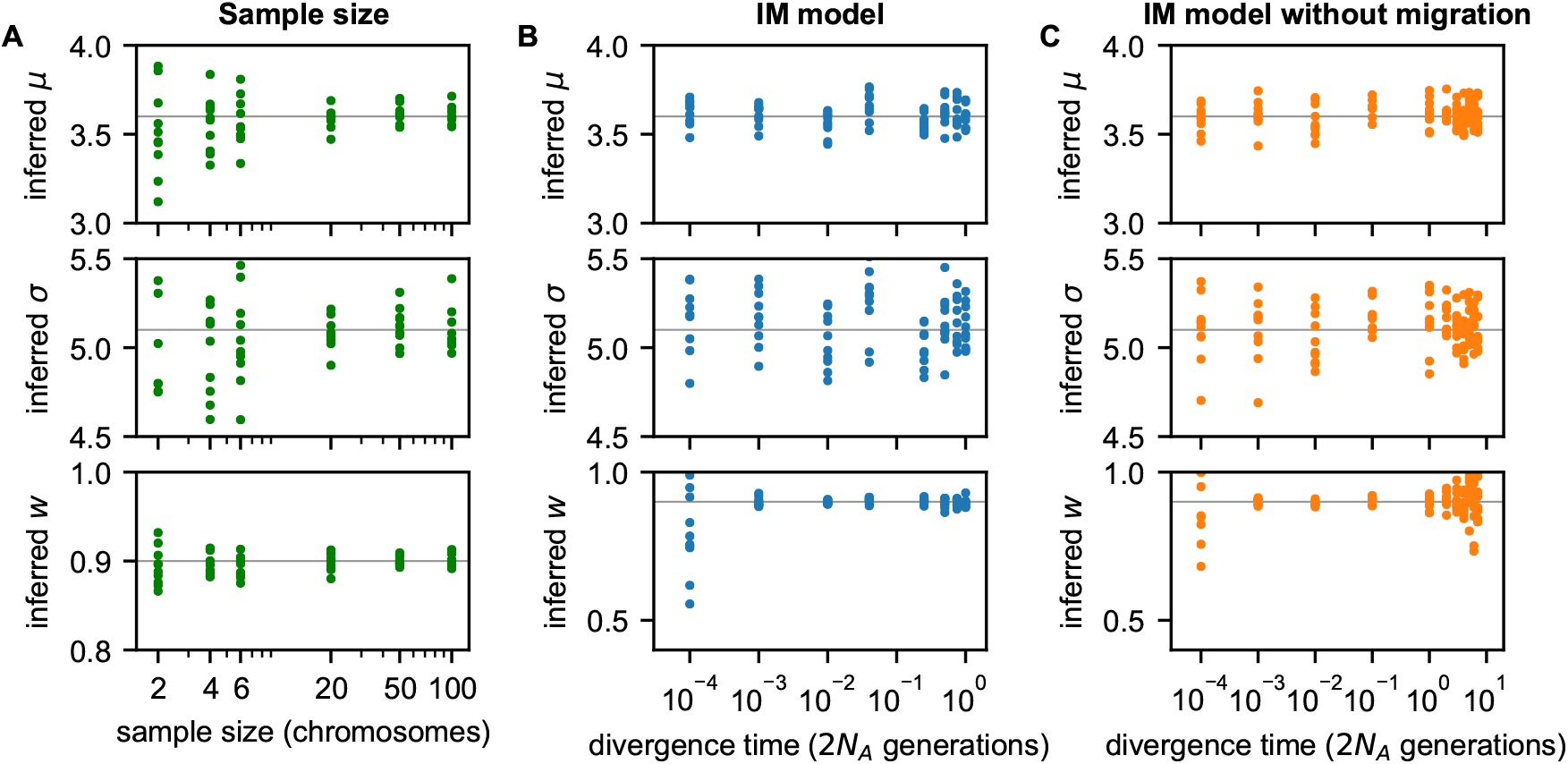
Precision of joint DFE inference versus sample sizes and divergence time. Simulated data were generated without linkage and with different sample sizes or divergence times, using the demographic model parameters in Table S1 with logornal *µ* = 3.6 and *σ* = 5.1 and DFE correlation *w* = 0.9. In each panel, points represent inferences from individual data sets and the gray line indicates the true value. **A**: Inferred DFE parameters for different sample sizes. For small sample sizes, *w* is still inferred precisely, even though *µ* and *σ* are highly uncertain. **B**: Inferred DFE parameters for different divergence times, in a model with migration. Precision of *w* inference is low for small divergence times. **C**: Inferred DFE parameters for different divergence times, in a model without migration. In this case, precision of *w* inference is low for both small and large divergence times.

**Figure S4:**
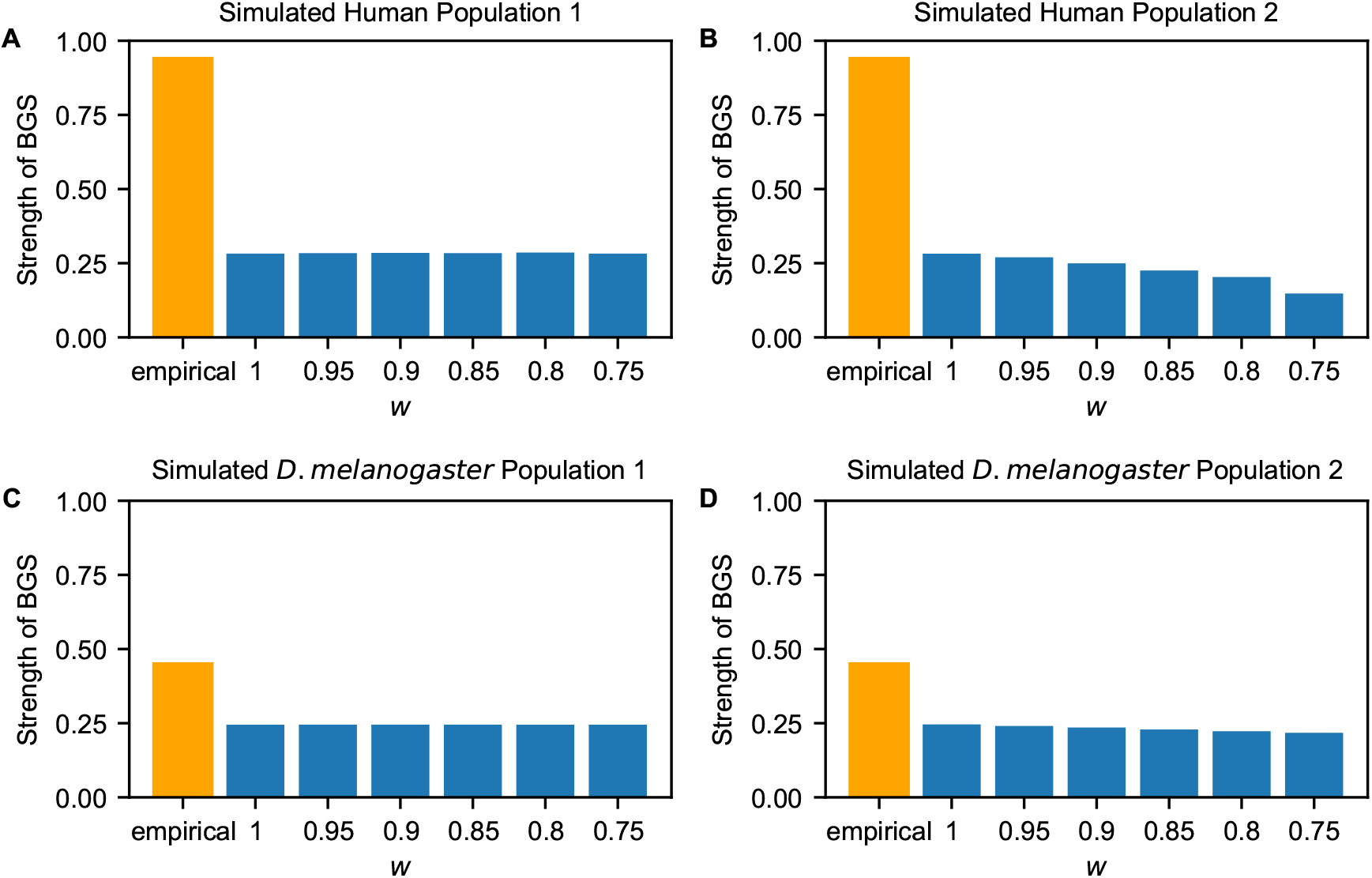
Strength of background selection (BGS) between simulation and empirical studies. The strengths of BGS in empirical studies are from Charlesworth (2013). The strengths in simulation are estimated from the observed number of pairwise differences between two chromosomes in the non-neutral scenarios versus the neutral ones. The data plotted in these figures can be found in Table S8.

**Figure S5:**
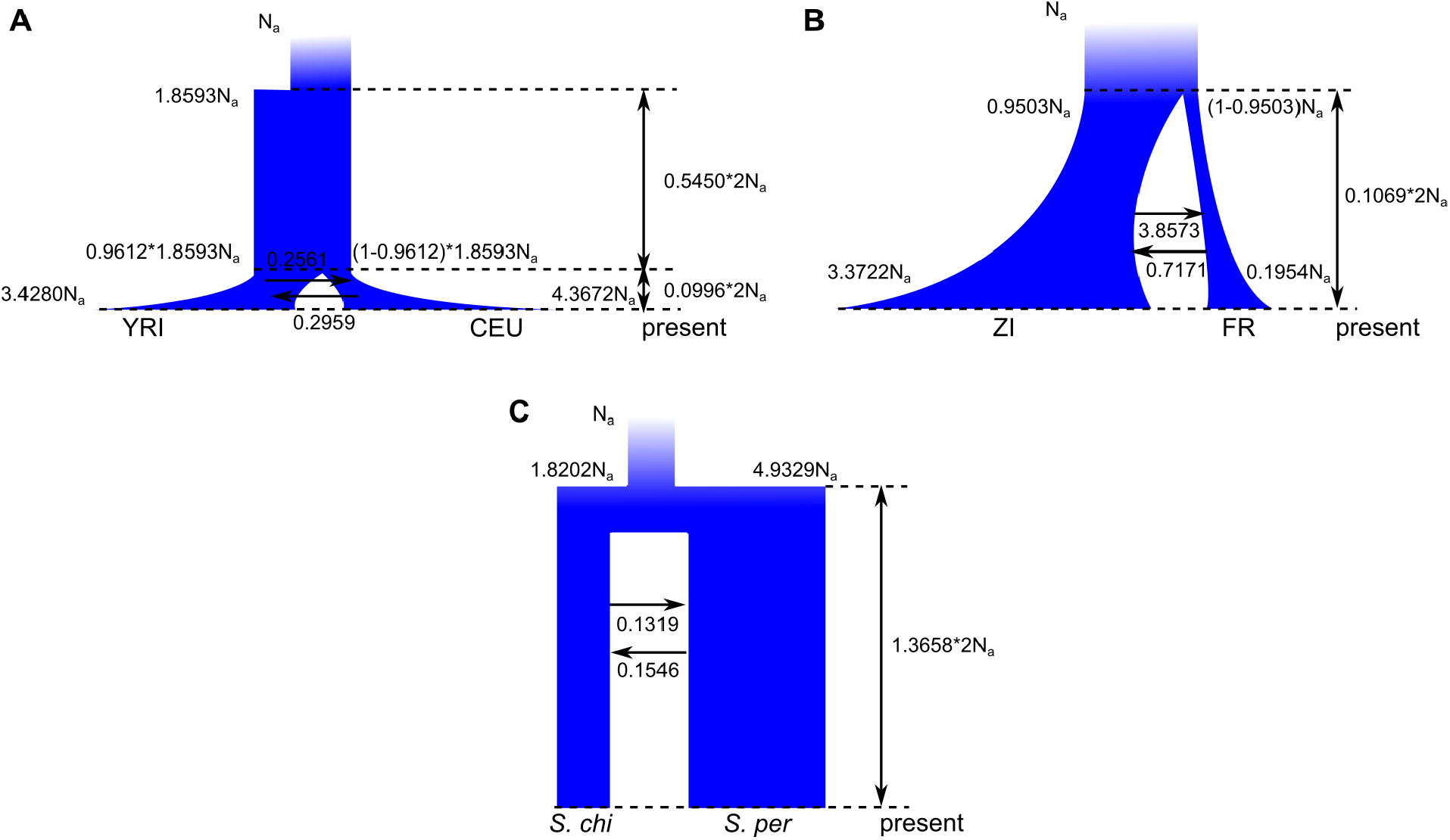
Best fit demographic models. **A**: The best fit demographic model for humans. **B**: The best fit demographic model for *D. melanogaster*. **C**: The best fit demographic model for wild tomatoes.

**Figure S6:**
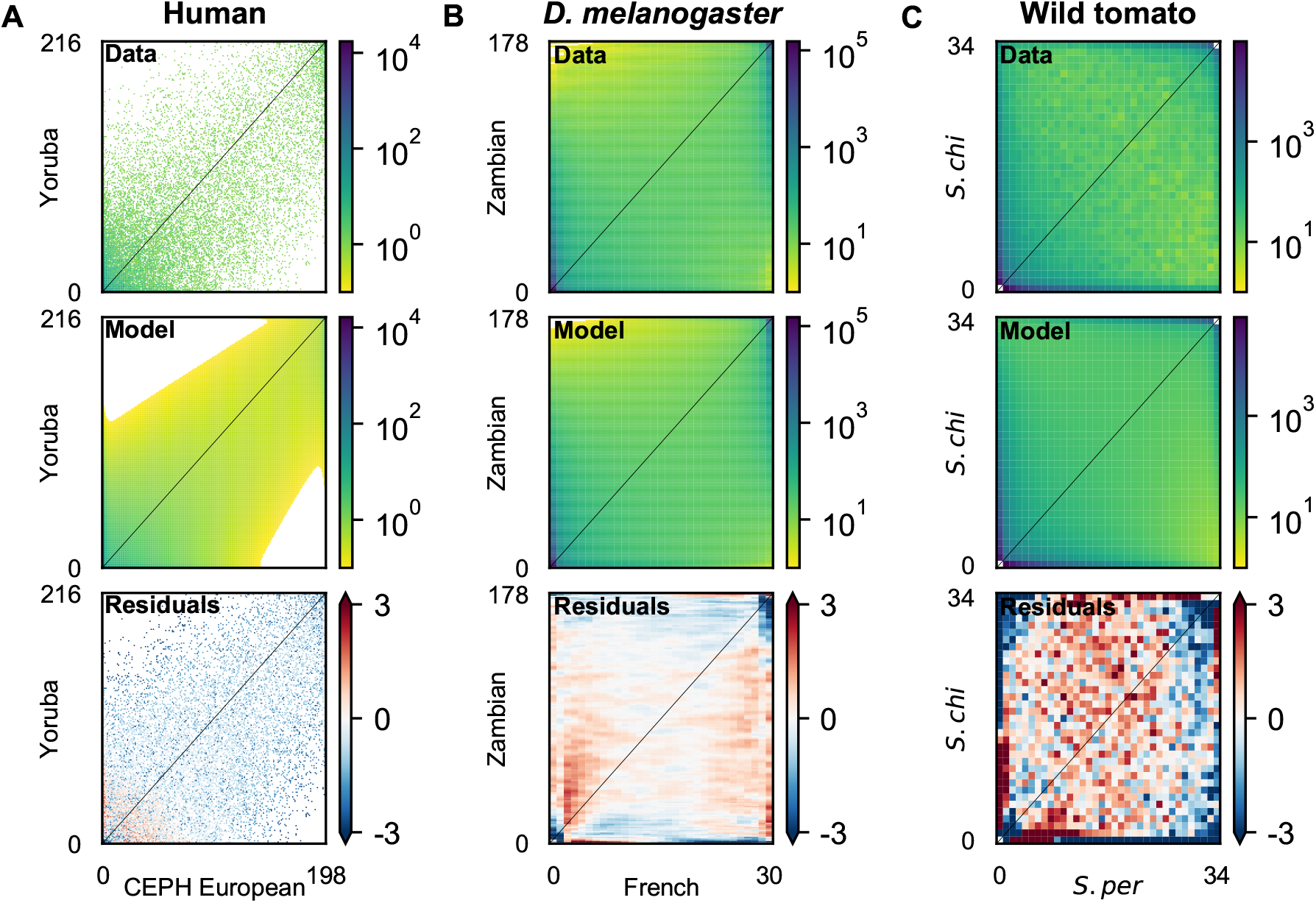
Model fits to joint allele frequency spectra (AFS) using synonymous data. **A**: Joint AFS for the human synonymous data and the best fit model (Fig. S5A). **B**: Joint AFS for the *D. melanogaster* synonymous data and the best fit model (Fig. S5B). **C**: Joint AFS for the wild tomato synonymous data and the best fit model (Fig. S5C).

**Figure S7:**
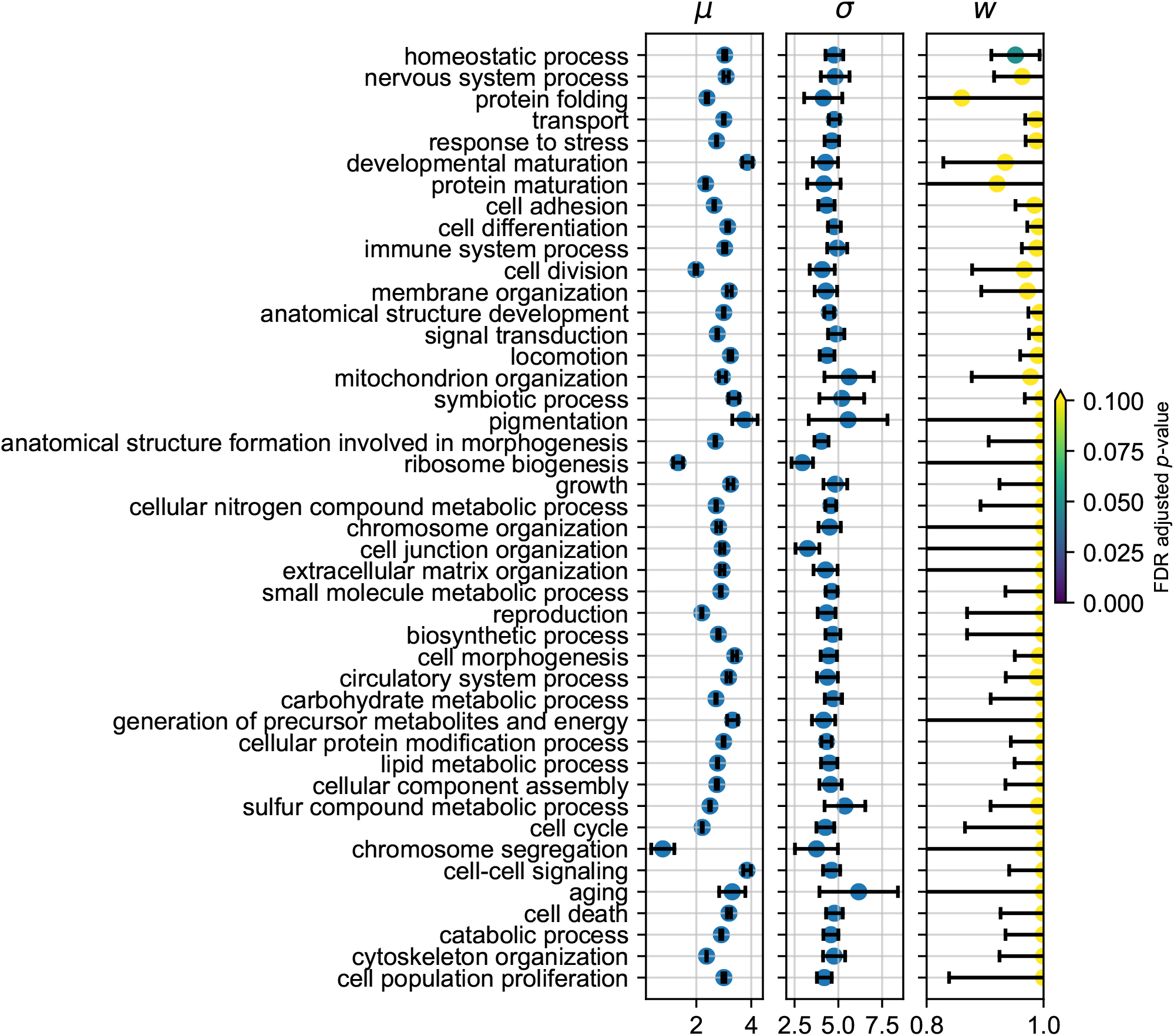
Joint DFE inference for different GO terms in humans. Plotted are maximum likelihood inferences with 95% confidence intervals. For inferred *w*, colors indicate FDR-adjusted *p*-values from two-tailed *z*-tests as to whether the confidence interval overlaps *w* = 1. The data plotted in these figures can be found in Table S9.

**Figure S8:**
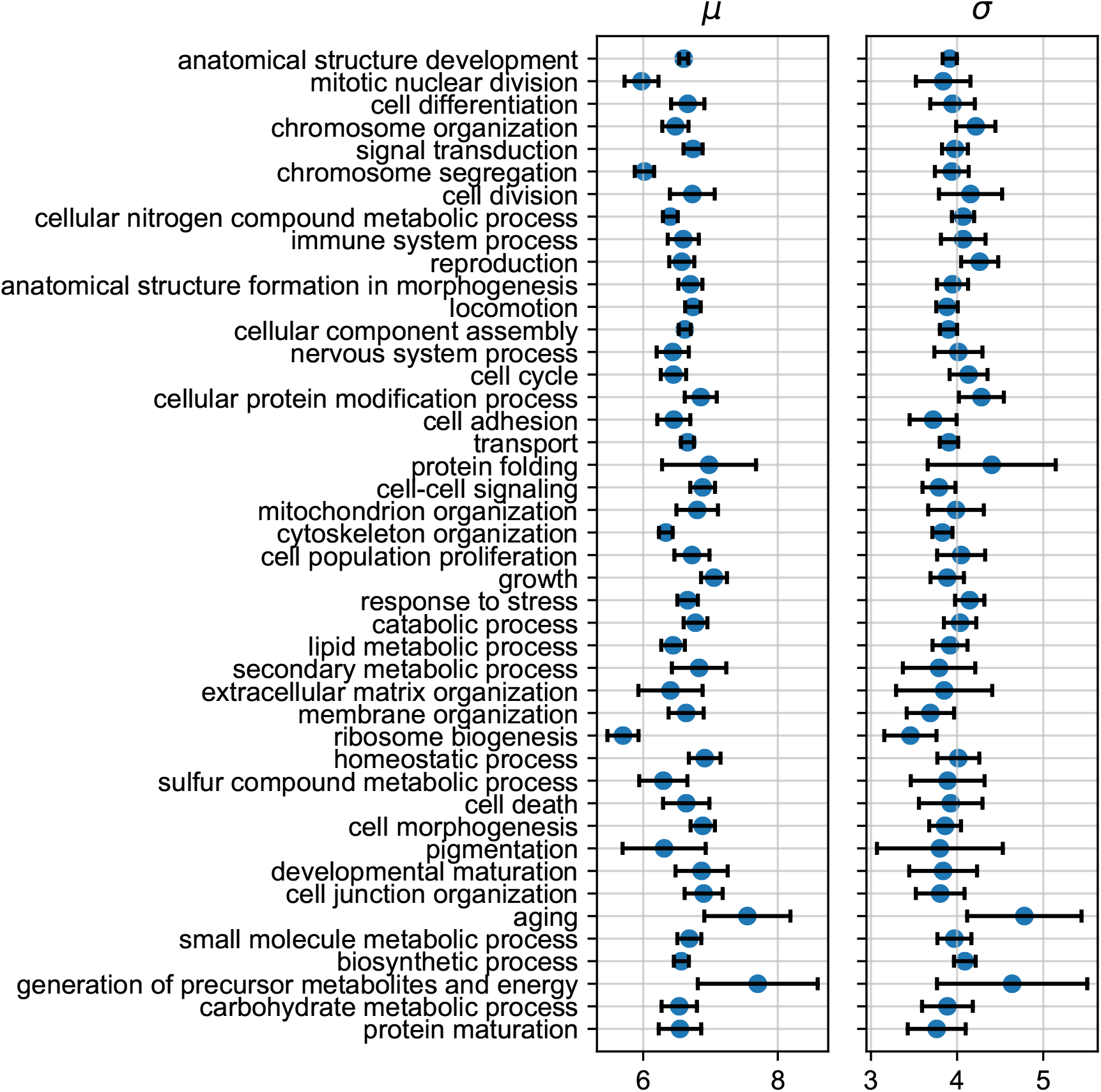
Inferred *µ* and *σ* from the joint DFE inference for different GO terms in *D. melanogaster*. Plotted are maximum likelihood inferences with 95% confidence intervals. The data plotted in these figures can be found in Table S10.

**Figure S9:**
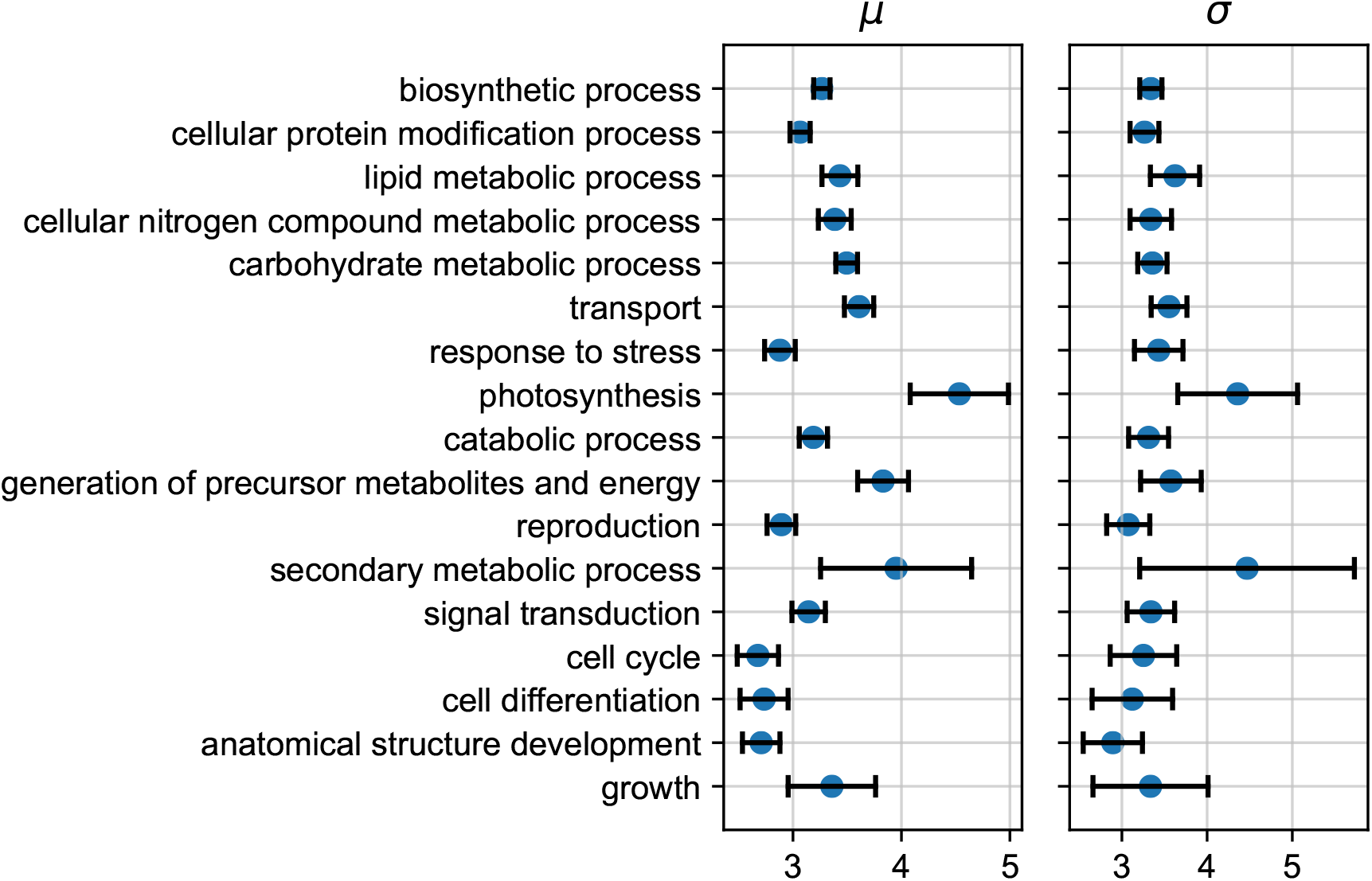
Inferred *µ* and *σ* from the joint DFE inference for different GO terms in wild tomatoes. Plotted are maximum likelihood inferences with 95% confidence intervals. The data plotted in these figures can be found in Table S11.

**Figure S10:**
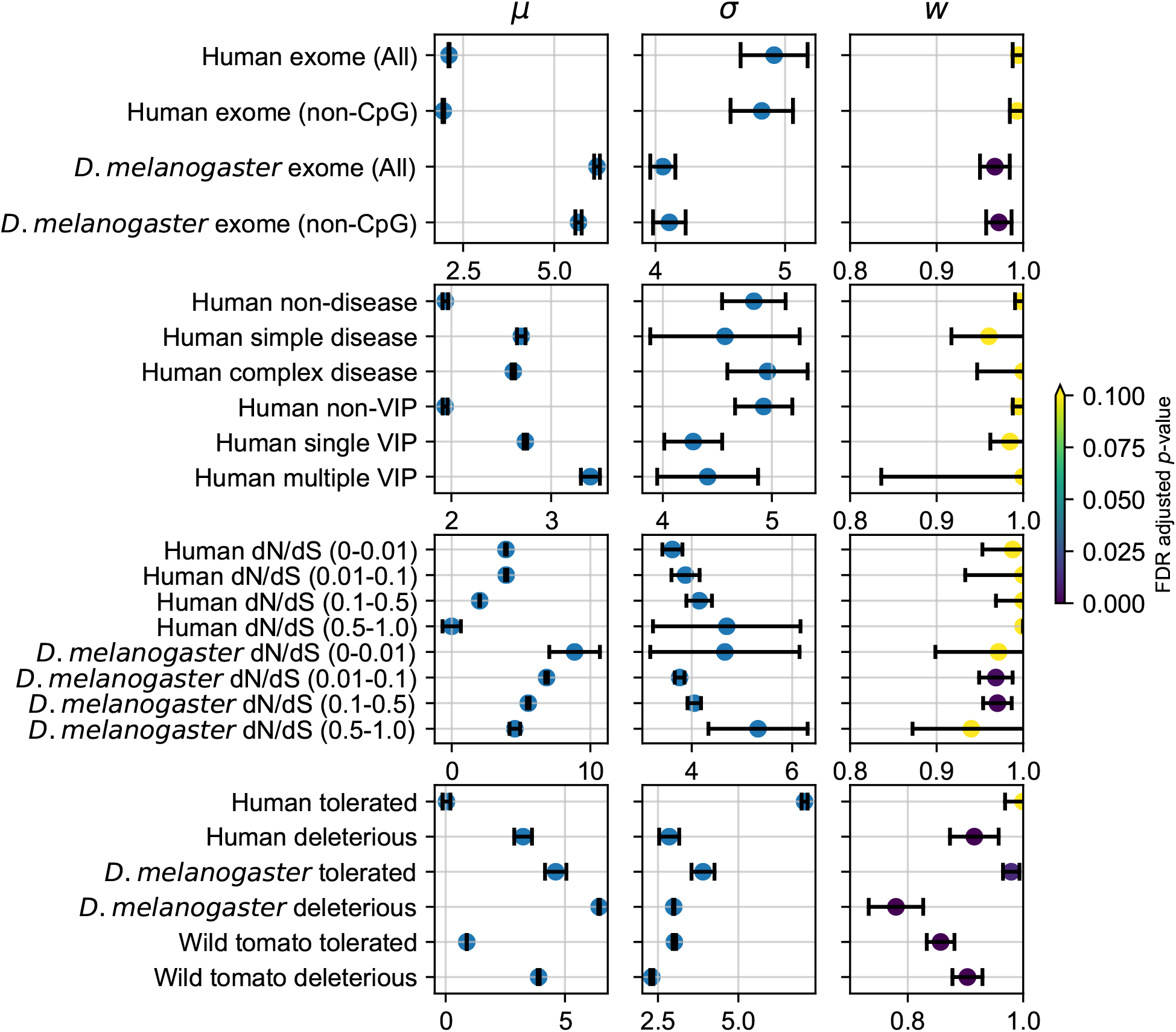
Joint DFE inference for different types of data in humans and *D. melanogaster*. Plotted are maximum likelihood inferences with 95% confidence intervals. For inferred *w*, colors indicate FDR-adjusted *p*-values from two-tailed *z*-tests as to whether the confidence interval overlaps *w* = 1. The data plotted in these figures can be found in Table S9, S10 and S11.

**Figure S11:**
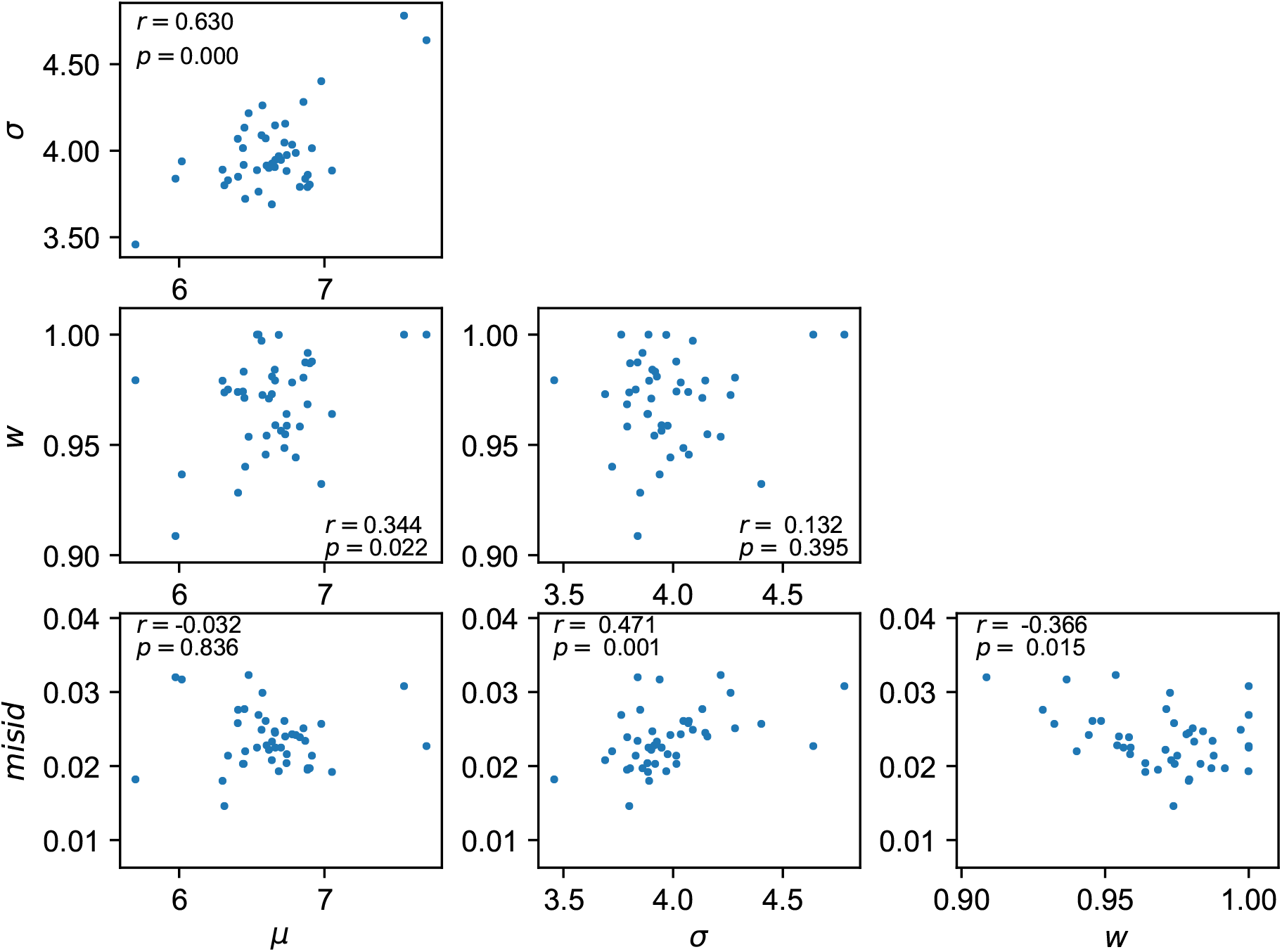
Relationships among among fitted parameters to *D. melanogaster* GO terms. Insets indicate Pearson correlations. The data plotted in these figures can be found in Table S10.

**Figure S12:**
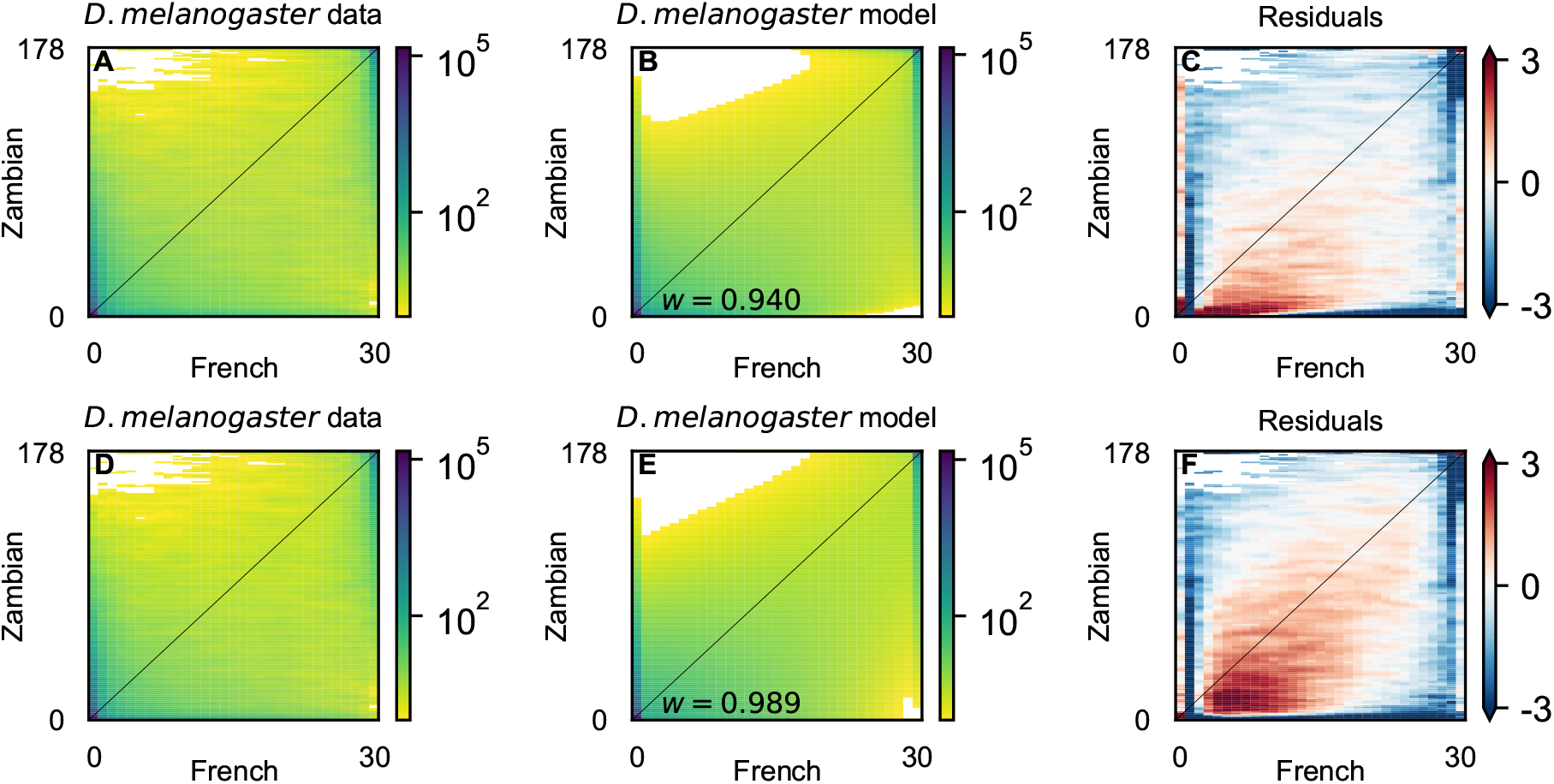
*D. melanogaster* results assuming simpler demographic models. **A, B & C**: A split mig model (Fig. S2C) with symmetric migration fits the data less well than our full model with exponential growth and asymmetric migration. (Compare residuals with Fig. S6F.) **D, E & F**: A split mig model without migration fits even worse.

**Figure S13:**
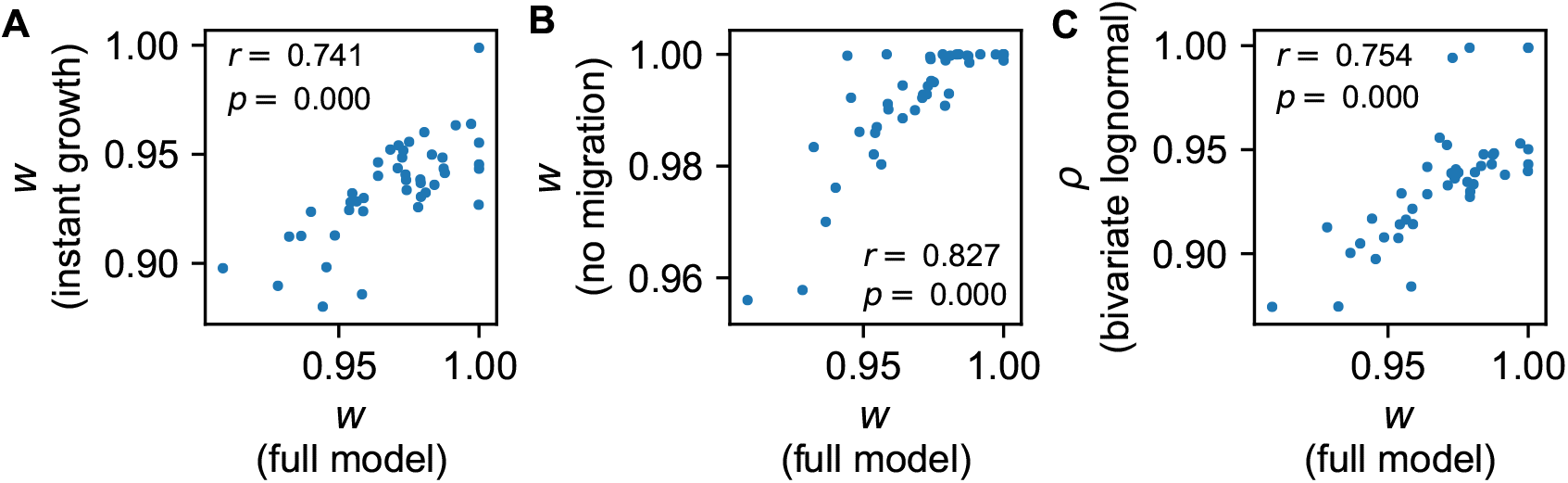
*D. melanogaster* results assuming different models. For the *D. melanogaster* Gene Ontology terms, inferences from the full demographic model and lognormal mixture DFE model are compared with **A**: A split mig model with symmetric migration, **B**: A split mig model withnot migration and **C**: A DFE model with a bivariate lognormal distribution. The data plotted in these figures can be found in Table S12.

**Figure S14:**
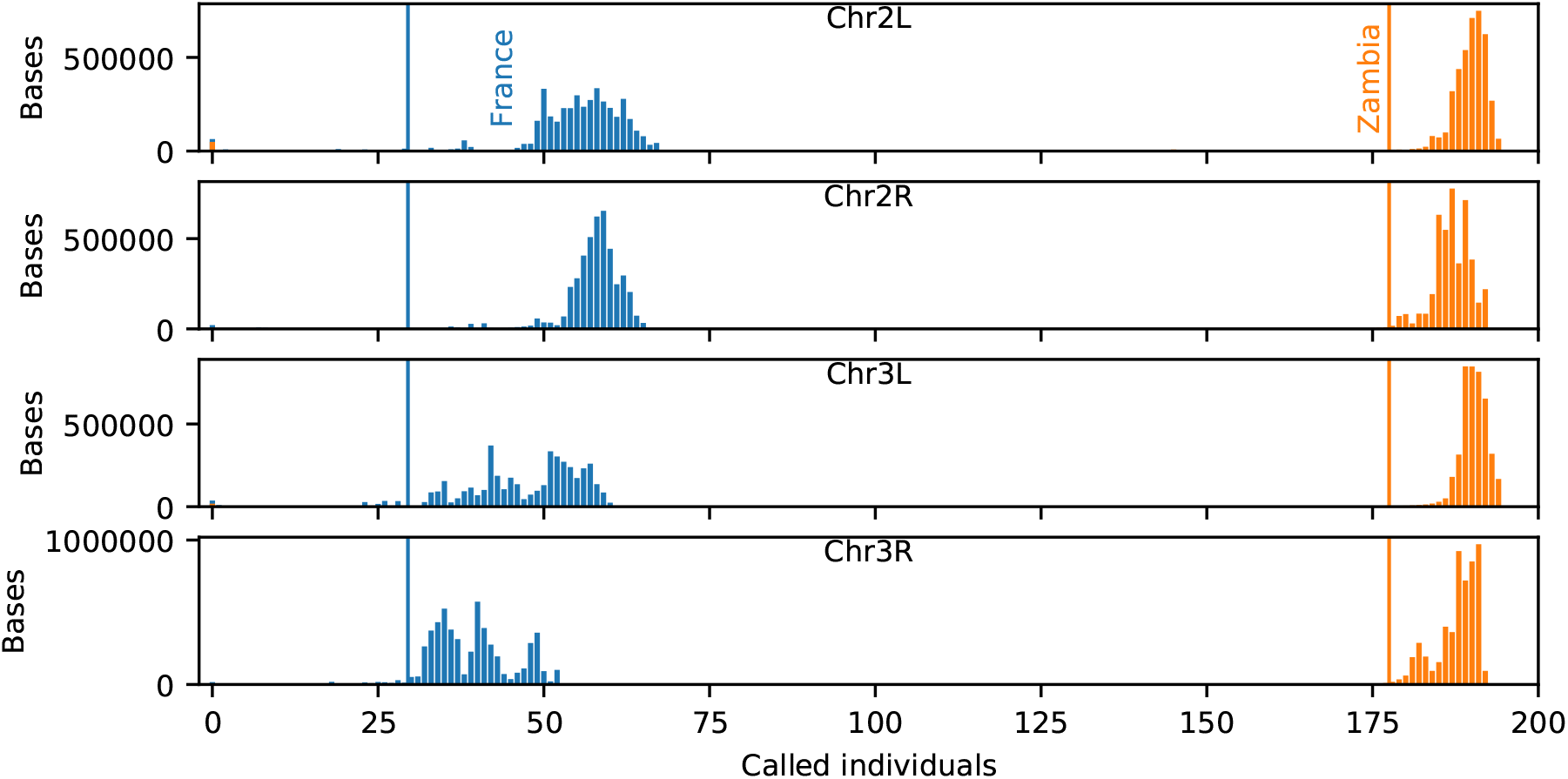
Calling rate for different chromosome arms in analyzed *D. melanogaster* data. For each arm, histograms indicate the number of bases at which a given number of individuals were called in each population. The vertical lines indicate the projection sizes for the AFS analysis.

**Figure S15:**
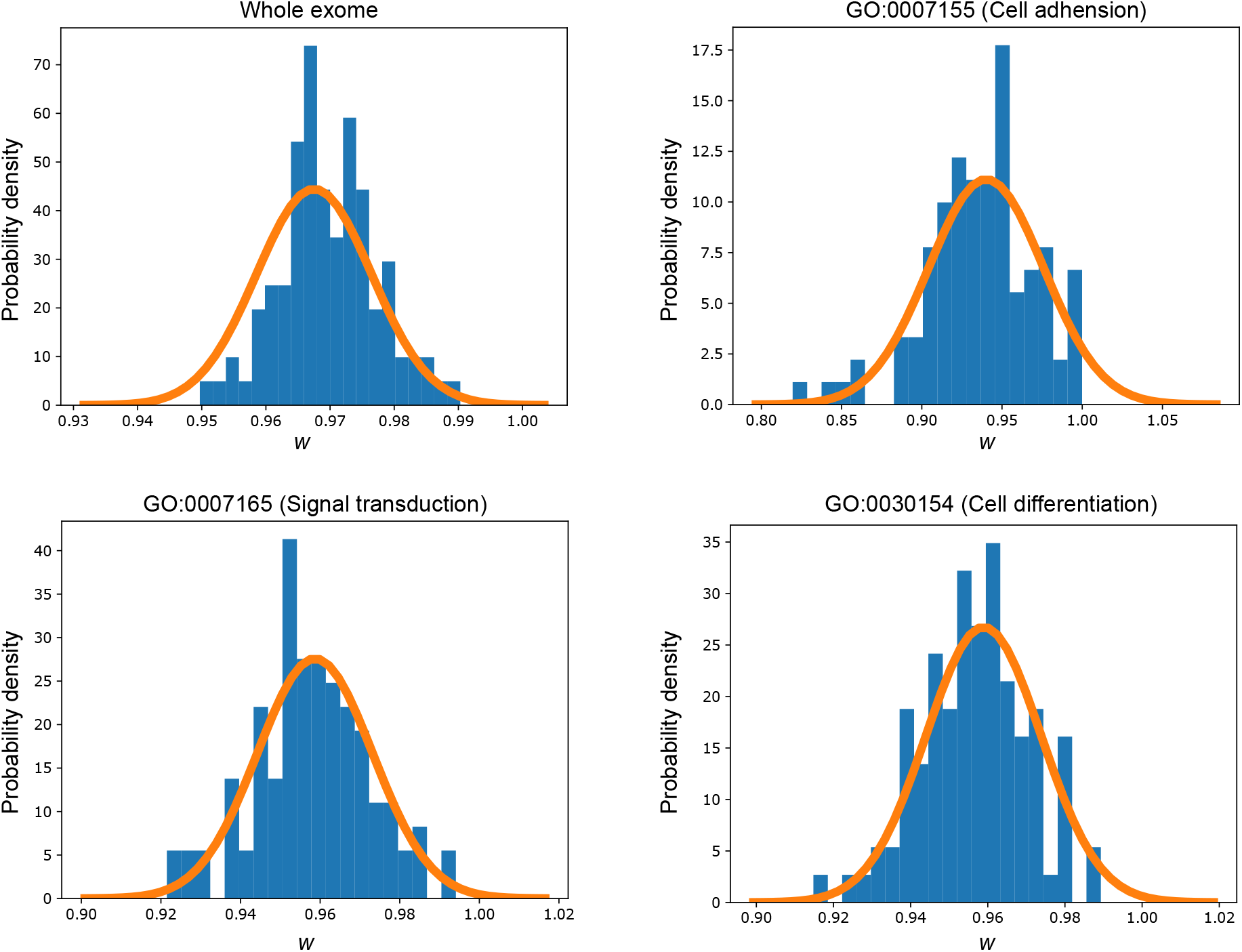
Comparison between uncertainties estimated with the Godambe Information Matrix and bootstrap fitting. For the *D. melanogaster* data, each panel shows a different subset of genes/mutations. In each panel, the histogram shows results from conventional bootstrap fitting, while the smooth curve is a normal distribution centered at the maximum likelihood inferred value and standard deviation estimated using the Godambe approach.

